## Supplementary Information for "Lineage EM Algorithm for Inferring Latent States from Cellular Lineage Trees"

So Nakashima et al.

#### S1 Previous attempts of latent variable estimation from linage trees

The latent-variable estimation from sequantial datais originally used in audio recognition [1, 2] and natural language processing [3]. These methods are applicable to analyze the lineages obtained by the mother machine [4]. Researchers have tried to extend the latent-variable estimation from sequential to branching data like lineage trees under specific assumptions over the data. For example, the latent-variable estimation based on the kin-correlation was proposed by assuming that the state transition of cells satisfies the detailed balanced condition [5, 6]. Another work employed an algebraic invariance of the lineage tree under the condition that the latent variable is observable [7, 8]. Monte Carlo Markov Chain method for a specific hierarchical model was developed to estimate the dynamics of the transcriptional activity of mammalian cells [9]. Furthermore, clustering algorithms [10, 11], Monte-Carlo algorithms [12], and model selection [13] are proposed for lineage trees whose shapes are deterministic or prefixed. Some of the above methods [5, 6, 10] additionally assume that the states $\boldsymbol{x}$ of the cells at the end of lineage are available.

#### S2 Model setup and time slice formulation

We introduce the setup of our population dynamics, which is equivalent to that given in Section 2 in the main text. Let $a \in [0, \infty)$ and $\boldsymbol{x} \in \Omega$ be an age and a state of cells, respectively, where $\Omega$ is either discrete or continuous state set of $\boldsymbol{x}$. Here, the age $a$ means the elapsed time since the last division. In the time evolution, a cell divides into two daughter cells with an age- and state-dependent rate $r_{\mathrm{F}}(a, \boldsymbol{x}) \geq 0$. With this rate, the probability density distribution of the division time can be represented as

$$\pi_{\mathrm{F}}(\tau|\boldsymbol{x}) := r_{\mathrm{F}}(\tau, \boldsymbol{x})\, e^{-\int_0^\tau r_{\mathrm{F}}(a,\boldsymbol{x})da}. \tag{S2.1}$$

where $e^{-\int_0^\tau r_{\mathrm{F}}(a,\boldsymbol{x})da}$ is the probability that the cell did not divide up to time $\tau$ and $r_{\mathrm{F}}(\tau, \boldsymbol{x})\, d\tau$ is the probability that the cell divides at age $\tau$. Here, we note that $\pi_{\mathrm{F}}(\tau|\boldsymbol{x})$ can be expressed as Eq. (S2.1) without loss of generality; that is, if we firstly obtain $\pi_{\mathrm{F}}(\tau|\boldsymbol{x})$, then we can calculate the division rate $r_{\mathrm{F}}(a, \boldsymbol{x})$ as

$$r_{\mathrm{F}}(a, \boldsymbol{x}) = \frac{\pi_{\mathrm{F}}(a|\boldsymbol{x})}{\int_a^\infty \pi_{\mathrm{F}}(\tau|\boldsymbol{x})\, d\tau}. \tag{S2.2}$$

Next, we define the probability of the state-switching from $\boldsymbol{x}'$ to $\boldsymbol{x}$ upon division by the transition matrix $\mathbb{T}_{\mathrm{F}}(\boldsymbol{x}|\boldsymbol{x}')$, where the state-switching of a daughter cell is dependent only on the state of the mother cell $\boldsymbol{x}'$ but independent of the state of its sister. We assume that $\mathbb{T}_{\mathrm{F}}$ is primitive (ergodic): there exists $n \in \mathbb{N}$ such that $\mathbb{T}_{\mathrm{F}}^n(\boldsymbol{x}'|\boldsymbol{x}) > 0$ for all $\boldsymbol{x}, \boldsymbol{x}' \in \Omega$.

Under this setup, the time evolution of the expected number of cells with the age $a$ and the state $\boldsymbol{x}$ at time $t$, which we denote by $N_t(a, \boldsymbol{x})$, can be described by the McKendric equation:

$$\frac{\partial}{\partial t} N_t(a, \boldsymbol{x}) = \left[-\frac{\partial}{\partial a} - r_{\mathrm{F}}(a, \boldsymbol{x})\right] N_t(a, \boldsymbol{x}), \tag{S2.3}$$

with a boundary condition,

$$N_t(0, \boldsymbol{x}) = 2 \sum_{x' \in \Omega} \int_0^\infty d\tau'\, \mathbb{T}_{\mathrm{F}}(\boldsymbol{x}|\boldsymbol{x}')\, r_{\mathrm{F}}(\tau', \boldsymbol{x}')\, N_t(\tau', \boldsymbol{x}'). \tag{S2.4}$$

The derivation of this equation is shown in Ref. [14]. By solving Eq. (S2.3) with the boundary condition Eq. (S2.4) , we obatin the time-slice distribution of the population $N_t(a, \boldsymbol{x}; a_0, \boldsymbol{x}_1)$, where $(a_0, \boldsymbol{x}_1)$ stands for the initial condition $N_0(a, \boldsymbol{x}) = \delta(a - a_0)\,\delta_{x,\boldsymbol{x}_1}$. Here, $\delta(\cdot)$ is the Dirac's delta function and $\delta_{\boldsymbol{x},\boldsymbol{x}'}$ is the Kronecker's delta.

### S3 Brief explanation for the chronological sampling

In the following section, we briefly introduce a pathwise formulation without mathematical rigorousness. More ditailed derivation and explanation for that are shown in Ref. [14]. In contrast to the above formation with partial differential equations, we here introduce a path-wise formulation. First, we consider the chronological sampling, where we time-forwardly track a dividing single cell by ignoring one of daughter cells. Suppose that we observe a path $\chi_T = \{n; a_0, (\tau_1, \boldsymbol{x}_1), ..., (\tau_n, \boldsymbol{x}_n), (\tau_{n+1}, \boldsymbol{x}_{n+1})\}$ in the chronological tracking up to time $T$, where $n$ is the number of division events during the observation, $\tau_i$ and $\boldsymbol{x}_i$ are the division time and the state of the cell before the $i$th division, and $a_0$ is the age of the root cell at time 0. For notational simplicity, $\tau_{n+1}$ is specially defined as the time interval between $n$th division and the time $T$. The probability $\mathbb{P}_{\mathrm{F}}[\chi_T]$ to observe such a path is calculated as

$$\mathbb{P}_{\mathrm{F}}[\chi_T] = \delta\left(T - \left\{\sum_{i=1}^{n+1}\tau_i - a_0\right\}\right) e^{-\int_0^{\tau_{n+1}} r_{\mathrm{F}}(a,\boldsymbol{x}_{n+1})da} \left[\prod_{i=1}^{n}\mathbb{T}_{\mathrm{F}}(\boldsymbol{x}_{i+1}|\boldsymbol{x}_i)\,\pi_{\mathrm{F}}(\tau_i|\boldsymbol{x}_i)\right] e^{\int_0^{a_0} r_{\mathrm{F}}(a,\boldsymbol{x}_1)da}, \tag{S3.1}$$

because it can be represented by the iteration of the the distribution of the division time $\pi_{\mathrm{F}}(\tau|\boldsymbol{x})$ and the transition matrix $\mathbb{T}_{\mathrm{F}}(\boldsymbol{x}|\boldsymbol{x}')$. Also, we used the fact that the probability until the 1st division can be represented as

$$r_{\mathrm{F}}(\tau_1, \boldsymbol{x}_1)\, e^{-\int_{a_0}^{\tau_1} r_{\mathrm{F}}(a,\boldsymbol{x}_1)da} = \pi_{\mathrm{F}}(\tau_1|\boldsymbol{x}_1)\, e^{\int_0^{a_0} r_{\mathrm{F}}(a,\boldsymbol{x}_1)da}. \tag{S3.2}$$

Here, we note that the path probability $\mathbb{P}_{\mathrm{F}}[\chi_T]$ can be regarded as the semi-Markov process (Markov renewal process) with the kernel $Q_{\mathrm{F}}(\boldsymbol{x}, \tau'|\boldsymbol{x}') := \mathbb{T}_{\mathrm{F}}(\boldsymbol{x}|\boldsymbol{x}')\,\pi_{\mathrm{F}}(\tau'|\boldsymbol{x}')$. Therefore, in the chronological sampling for a sufficiently long time, the empirical histograms $\pi_{\mathrm{emp}}$ and $\mathbb{T}_{\mathrm{emp}}$ (Eqs. [1] and [2] in the main text) converge to the generators of the semi-Markov process $\pi_{\mathrm{F}}$ and $\mathbb{T}_{\mathrm{F}}$, respectively.

### S4 Brief explanation for the retrospective sampling

Next, we formulate the retrospective process and consider the convergence property of the empirical histograms in the retrospective sampling. Let $N_T[\chi_T]$ be the expected number of cells at time $T$ that have a history (path) $\chi_T$. On the cell-lineage tree $\mathcal{T}$ up to time $T$, $N_T[\chi_T]$ can be interpreted as the expected number of the paths that penetrate from time 0 to time $T$ in $\mathcal{T}$ with the history $\chi_T$. This number $N_T[\chi_T]$ can be evaluated as

$$N_T[\chi_T] = 2^n \mathbb{P}_{\mathrm{F}}[\chi_T], \tag{S4.1}$$

because the population doubles upon each division event. Details for a derivation of Eq. (S4.1) are shown in Appendix C of Ref. [14]. By dividing Eq. (S4.1) by the total number of the cells at time $T$, we define the following path probability:

$$\mathbb{P}_{\mathrm{Back}}[\chi_T] := \frac{N_T[\chi_T]}{N_T^{\mathrm{tot}}}, \tag{S4.2}$$

where the total population $N_T^{\mathrm{tot}}$ is calculated as

$$N_T^{\mathrm{tot}} = \sum_{n=0}^{\infty}\sum_{\{\boldsymbol{x}_i\}_{i=0}^{n}}\int_0^{\infty}\cdots\int_0^{\infty}\prod_{i=1}^{n} d\tau_i\, N_T[\chi_T]. \tag{S4.3}$$

$\mathbb{P}_{\mathrm{Back}}[\chi_T]$ represents the probability to observe $\chi_T$ when we uniformly sample a cell at time $T$ from a cultured population and retrospectively trace its ancestors back, which means that the path probability $\mathbb{P}_{\mathrm{Back}}[\chi_T]$ is obtained in the retrospective sampling. Furthermore, $\mathbb{P}_{\mathrm{Back}}[\chi_T]$ can be approximated by the semi-Markov process generated by $\pi_{\mathrm{B}}$ and $\mathbb{T}_{\mathrm{B}}$ in the main text (see Eqs. [3] and [4]). To clarify that, we substitute Eqs. [3] and [4] into Eq. (S4.2). Then, we obtain

$$\mathbb{P}_{\mathrm{Back}}[\chi_T] = \delta\left(T - \left[\sum_{i=1}^{n+1} \tau_i - a_0\right]\right) e^{-\lambda \tau_{n+1}} \frac{e^{\lambda T}}{N_{\mathrm{tot}}}$$

$$\frac{u(\boldsymbol{x}_1) e^{-\int_0^{\tau_{n+1}} r_F(a,\boldsymbol{x}_{n+1}) da}}{u(\boldsymbol{x}_{n+1}) e^{-\int_0^{a_0} r_F(a,\boldsymbol{x}_1) da}} \prod_{i=1}^{n} \mathbb{T}_{\mathrm{B}}(\boldsymbol{x}_{i+1}|\boldsymbol{x}_i) \pi_{\mathrm{B}}(\tau_i|\boldsymbol{x}_i)$$

$$= e^{-\lambda \tau_{n+1}} \frac{e^{\lambda T}}{N_{\mathrm{tot}}} \cdot \frac{u(\boldsymbol{x}_1) e^{-\int_0^{\tau_{n+1}} r_F(a,\boldsymbol{x}_{n+1}) da} e^{-\int_0^{a_0} r_B(a,\boldsymbol{x}_1) da}}{u(\boldsymbol{x}_{n+1}) e^{-\int_0^{a_0} r_F(a,\boldsymbol{x}_1) da} e^{-\int_0^{\tau_{n+1}} r_B(a,\boldsymbol{x}_{n+1}) da}} \mathbb{P}_B[\chi_T], \tag{S4.4}$$

where $\mathbb{P}_B[\chi_T]$ represents the path probability of semi-Markov process with the kernel $Q_B(\boldsymbol{x}_{i+1}, \tau_i|\boldsymbol{x}_i) := \mathbb{T}_B(\boldsymbol{x}|\boldsymbol{x}') \pi_B(\tau'|\boldsymbol{x}')$:

$$\mathbb{P}_B[\chi_T] = \delta\left(T - \left\{\sum_{i=1}^{n+1} \tau_i - a_0\right\}\right) e^{-\int_0^{\tau_{n+1}} r_B(a,\boldsymbol{x}_{n+1}) da} \left[\prod_{i=1}^{n} Q_B(\boldsymbol{x}_{i+1}, \tau_i|\boldsymbol{x}_i)\right] e^{\int_0^{a_0} r_B(a,\boldsymbol{x}_1) da}. \tag{S4.5}$$

Here, $r_B(a,x)$ denotes the division rate defined as

$$r_B(a,\boldsymbol{x}) = \frac{\pi_{\mathrm{B}}(a|\boldsymbol{x})}{\int_a^\infty da' \, \pi_{\mathrm{B}}(a'|\boldsymbol{x})}. \tag{S4.6}$$

By neglecting boundary factors in Eq. (S4.4) and approximating $e^{\lambda T} \approx N_{\mathrm{tot}}$, we obtain $\mathbb{P}_B[\chi_T] \approx \mathbb{P}_{\mathrm{Back}}[\chi_T]$. Therefore, the backward process is mimicked by the semi-Markov process $\mathbb{P}_B[\chi_T]$ up to the boundary factors, which is called the retrospective process [14]. By using this result, we find that the empirical distributions (Eqs. [1] and [2] in the main text) for the retrospective sampling converge to the kernels of the semi-Markov process $\pi_B$ and $\mathbb{T}_B$ for a sufficiently long time such that we can igonore boundary factors. A more detailed proof is shown in Ref. [14].

Finally, we consider the connection of the pathwise formula (S4.4) to the time-slice distribution $N_T(a,\boldsymbol{x};a_0,\boldsymbol{x}_1)$, which is a solution of the McKendrick equation (S2.3). Since $N_T(a,\boldsymbol{x};a_0,\boldsymbol{x}_1)$ is the expected number of the cells at time $T$ with the age $a$ and the state $\boldsymbol{x}$ in the lineage tree, which corresponds to the leaf cells, we obtain

$$N_T(a,\boldsymbol{x};a_0,\boldsymbol{x}_1) = \sum_{n=0}^{\infty} \int d\tau_1 \prod_{i=2}^{n} \left[\sum_{\boldsymbol{x}_i \in \Omega} \int d\tau_i\right] \int d\tau_{n+1} \, \delta(a - \tau_{n+1}) \delta_{\boldsymbol{x},\boldsymbol{x}_{n+1}} N_T[\chi_T]. \tag{S4.7}$$

Here, $N_T[\chi_T]$ is calculated by Eqs. (S4.2) and (S4.4) as

$$N_T[\chi_T] = e^{\lambda(T-\tau_{n+1})} \frac{u(\boldsymbol{x}_1) e^{-\int_0^{\tau_{n+1}} r_F(a,\boldsymbol{x}_{n+1}) da} e^{-\int_0^{a_0} r_B(a,\boldsymbol{x}_1) da}}{u(\boldsymbol{x}_{n+1}) e^{-\int_0^{a_0} r_F(a,\boldsymbol{x}_1) da} e^{-\int_0^{\tau_{n+1}} r_B(a,\boldsymbol{x}_{n+1}) da}} \mathbb{P}_B[\chi_T]. \tag{S4.8}$$

Substituting this equation into (S4.7), we have

$$N_T(a,\boldsymbol{x};a_0,\boldsymbol{x}_1) = e^{\lambda(T-a)} \frac{u(\boldsymbol{x}_1) e^{-\int_0^{a} dt \, r_F(t,\boldsymbol{x})} e^{-\int_0^{a_0} dt \, r_B(t,\boldsymbol{x}_1)}}{u(\boldsymbol{x}) e^{-\int_0^{a_0} dt \, r_F(t,\boldsymbol{x}_1)} e^{-\int_0^{a} dt \, r_B(t,\boldsymbol{x})}} p_T^B(a,\boldsymbol{x};a_0,\boldsymbol{x}_1), \tag{S4.9}$$

where $p_T^B$ is the time-slice distribution of the retrospective process at time $T$ defined by

$$p_T^B(a,\boldsymbol{x};a_0,\boldsymbol{x}_1) = \int d\tau_1 \prod_{i=2}^{n} \int d\tau_{n+1} \, \delta(a - \tau_{n+1}) \left[\sum_{x_i \in \Omega} \int d\tau_i\right] \mathbb{P}_B[\chi_T]. \tag{S4.10}$$

This representation is known as many-to-one formula for leaves [15, 16].

#### S5 A sketch of the proof of the convergence of the empirical distributions for the tree sampling

In this section, we give a sketch of the proof that the empirical distributions (Eqs. [1] and [2] in the main text) converge to $\pi_{\mathrm{B}}$ and $\mathbb{T}_{\mathrm{F}}$ if we choose the set of the cells $\mathcal{D}$ as the all non-leaf cells in lineage tree. We give the rigorous proof in the next section. We first show the convergence of the empirical distribution of the division time, which is defined by

$$\pi_{\mathrm{emp}}^{\mathcal{T}_{\boldsymbol{x}}}(\tau|\boldsymbol{x}) = \frac{1}{|\mathcal{T}_{\boldsymbol{x}}|}\sum_{i\in\mathcal{T}_{\boldsymbol{x}}}\delta(\tau-\tau_i). \tag{S5.1}$$

Here, $\mathcal{T}_{\boldsymbol{x}}$ is the non-leaf cells with the state $\boldsymbol{x}$ in the lineage tree $\mathcal{T}$ and $|\mathcal{T}_{\boldsymbol{x}}|$ stands for the cardinality of the set $\mathcal{T}_{\boldsymbol{x}}$. Let $\mathring{N}_T(\tau,\boldsymbol{x};a_0,\boldsymbol{x}_1)$ be the expected number of the non-leaf cells in the lineage tree with the division time $\tau$ and the state $\boldsymbol{x}$ up to time $T$. Then, the empirical distribution of the division time (S5.1) is approximated as

$$\pi_{\mathrm{emp}}^{\mathcal{T}_{\boldsymbol{x}}}(\tau|\boldsymbol{x}) \approx \frac{\mathring{N}_T(\tau,\boldsymbol{x};a_0,\boldsymbol{x}_1)}{\mathring{N}_T^{\mathrm{tot}}(\boldsymbol{x})}, \tag{S5.2}$$

where $\mathring{N}_T^{\mathrm{tot}}(\boldsymbol{x})$ is the expected number of the non-leaf cells with the state $\boldsymbol{x}$ in the lineage tree: $\mathring{N}_T^{\mathrm{tot}}(\boldsymbol{x}) = \int d\tau \mathring{N}_T(\tau,\boldsymbol{x};a_0,\boldsymbol{x}_1)$. Here, we used two approximations: first, we assumed that $\pi_{\mathrm{emp}}^{\mathcal{T}_{\boldsymbol{x}}}(\tau|\boldsymbol{x})$ converges to its expectation value $\mathbb{E}\left[\left(\sum_{i\in\mathcal{T}_{\boldsymbol{x}}}\delta(\tau-\tau_i)\right)/|\mathcal{T}_{\boldsymbol{x}}|\right]$ as $T\to\infty$; second, we approximated this expectation as

$$\mathbb{E}\left[\frac{1}{|\mathcal{T}_{\boldsymbol{x}}|}\sum_{i\in\mathcal{T}_{\boldsymbol{x}}}\delta(\tau-\tau_i)\right] \approx \frac{\mathbb{E}\left[\sum_{i\in\mathcal{T}_{\boldsymbol{x}}}\delta(\tau-\tau_i)\right]}{\mathbb{E}\left[|\mathcal{T}_{\boldsymbol{x}}|\right]} = \frac{\mathring{N}_T(\tau,\boldsymbol{x};a_0,\boldsymbol{x}_1)}{\mathring{N}_T^{\mathrm{tot}}(\boldsymbol{x})}. \tag{S5.3}$$

In the next section, we give a rigorous proof without these approximations. In the following, we evaluate $\mathring{N}_T(\tau,\boldsymbol{x};a_0,\boldsymbol{x}_1)$. Let $D_t(\tau,\boldsymbol{x};a_0,\boldsymbol{x}_1)$ be the expected number of the cells that divide at time $t$ with the division time $\tau$ and the state $\boldsymbol{x}$. By integrating it with respect to $t$, we obtain

$$\mathring{N}_T(\tau,\boldsymbol{x};a_0,\boldsymbol{x}_1) = \int_0^T dt\, D_t(\tau,\boldsymbol{x};a_0,\boldsymbol{x}_1). \tag{S5.4}$$

By multiplying the solution of McKendrick equation (S4.7), $N_t(\tau,\boldsymbol{x};a_0,\boldsymbol{x}_1)$, by $r_F(\tau,\boldsymbol{x})$, we can calculate $D_t(\tau,\boldsymbol{x};a_0,\boldsymbol{x}_1)$ as

$$D_t(\tau,\boldsymbol{x};a_0,\boldsymbol{x}_1) = r_F(\tau,\boldsymbol{x})N_t(\tau,\boldsymbol{x};a_0,\boldsymbol{x}_1). \tag{S5.5}$$

For the rigorous derivation, see Lemma 1 in the next section. Equations (S4.9) , (S5.4), and (S5.5) lead to

$$\begin{aligned}&\mathring{N}_T(\tau,\boldsymbol{x};a_0,\boldsymbol{x}_1)\\ &= e^{-\lambda\tau}r_F(\tau,\boldsymbol{x})\frac{u(\boldsymbol{x}_1)e^{-\int_0^\tau da\, r_F(a,\boldsymbol{x})}e^{-\int_0^{a_0} da\, r_B(a,\boldsymbol{x}_1)}}{u(\boldsymbol{x})e^{-\int_0^{a_0} da\, r_F(a,\boldsymbol{x}_1)}e^{-\int_0^\tau da\, r_B(a,\boldsymbol{x})}}\int_0^T dt\, e^{\lambda t}p_t^B(\tau,\boldsymbol{x};a_0,\boldsymbol{x}_1).\end{aligned} \tag{S5.6}$$

Since, for sufficiently large $t$, $p_t^B(\tau,\boldsymbol{x};a_0,\boldsymbol{x}_1)$ converges to the stationary distribution of the semi-Markov process [17, 18] as

$$p_t^B(\tau,\boldsymbol{x};a_0,\boldsymbol{x}_1) \to p_{\mathrm{st}}^B(\tau,\boldsymbol{x}) = c(\boldsymbol{x})e^{-\int_0^\tau da\, r_B(a,\boldsymbol{x})}, \tag{S5.7}$$

where $c(\boldsymbol{x})$ is the stationary density of the cells with the age 0 and the state $\boldsymbol{x}$ of the retrospective process. Hence, we have

$$\int_0^T dt\, e^{\lambda t}p_t^B(\tau,\boldsymbol{x};a_0,\boldsymbol{x}_1) \approx c(\boldsymbol{x})e^{-\int_0^\tau da\, r_B(a,\boldsymbol{x})}\int_0^T dt\, e^{\lambda t} = c(\boldsymbol{x})e^{-\int_0^\tau da\, r_B(a,\boldsymbol{x})}\frac{e^{\lambda T}-1}{\lambda}. \tag{S5.8}$$

By substituting Eq. (S5.8) into Eq. (S5.6) and approximating $\mathring{N}_{\mathrm{tot}}(\boldsymbol{x})$ by $d(\boldsymbol{x})e^{\lambda T}$ for a certain state-dependent constant $d(\boldsymbol{x})$, we have

$$\begin{aligned}\pi_{\mathrm{emp}}^{\mathcal{T}_{\boldsymbol{x}}}(\tau|\boldsymbol{x}) &\approx e^{-\lambda\tau}r_F(\tau,\boldsymbol{x})\frac{u(\boldsymbol{x}_1)e^{-\int_0^\tau da\, r_F(a,\boldsymbol{x})}e^{-\int_0^{a_0} da\, r_B(a,\boldsymbol{x}_1)}}{u(\boldsymbol{x})e^{-\int_0^{a_0} da\, r_F(a,\boldsymbol{x}_1)}e^{-\int_0^\tau da\, r_B(a,\boldsymbol{x})}}c(\boldsymbol{x})e^{-\int_0^\tau da\, r_B(a,\boldsymbol{x})}\frac{e^{\lambda T}-1}{\lambda d(\boldsymbol{x})e^{\lambda T}}\\ &\propto e^{-\lambda\tau}\pi_{\mathrm{F}}(\tau|\boldsymbol{x}).\end{aligned} \tag{S5.9}$$

In the second line in Eq. (S5.9), we focus on the factors relating to $\tau$ and use Eq. (S2.1). Hence, we find that $\pi^{\mathcal{T}_{\boldsymbol{x}}}_{\mathrm{emp}}(\tau|\boldsymbol{x})$ converges to $\pi_{\mathrm{B}}(\tau|\boldsymbol{x})$:

$$\pi_{\mathrm{B}}(\tau|\boldsymbol{x}) = \frac{e^{-\lambda\tau}\pi_{\mathrm{F}}(\tau|\boldsymbol{x})}{Z(\boldsymbol{x})}, \tag{S5.10}$$

where $Z(\boldsymbol{x})$ is the normalization constant of $e^{-\lambda\tau}\pi_{\mathrm{F}}(\tau|\boldsymbol{x})$.

Next, we consider the convergence property of the empirical distribution of the state-switching Eq. [4] in the main text, which is almost the same procedure as the above derivation. Recall that the empirical distribution of the state transition is defined by

$$\mathbb{T}^{\mathcal{T}_{\boldsymbol{x}}}_{\mathrm{emp}}(\boldsymbol{x}'|\boldsymbol{x}) = \frac{\text{the number of the transitions from } \boldsymbol{x} \text{ to } \boldsymbol{x}'}{\text{the number of the transitions from } \boldsymbol{x}}. \tag{S5.11}$$

Let $\mathring{N}_T\left(\boldsymbol{x}' \leftarrow \boldsymbol{x}; a_0, x_1\right)$ be the expected number of the state-switching from state $\boldsymbol{x}$ to state $\boldsymbol{x}'$ in a lineage tree up to time $T$. Then, the empirical distribution of the state-switching (S5.11) can be approximated as

$$\mathbb{T}^{\mathcal{T}_{\boldsymbol{x}}}_{\mathrm{emp}}(\boldsymbol{x}'|\boldsymbol{x}) \approx \frac{\mathring{N}_T\left(\boldsymbol{x}' \leftarrow \boldsymbol{x}; a_0, \boldsymbol{x}_1\right)}{2\mathring{N}^{\mathrm{tot}}_T(\boldsymbol{x})}, \tag{S5.12}$$

where the prefactor 2 stands for the number of the daughters. In the following, we evaluate $\mathring{N}_T\left(\boldsymbol{x}' \leftarrow \boldsymbol{x}; a_0, \boldsymbol{x}_1\right)$. Let $D_t\left(\boldsymbol{x}' \leftarrow \boldsymbol{x}; a_0, \boldsymbol{x}_1\right)$ be the expected number of the state-switching from state $\boldsymbol{x}$ to state $\boldsymbol{x}'$ at time $t$. By integration, we obtain

$$\mathring{N}_T\left(\boldsymbol{x}' \leftarrow \boldsymbol{x}; a_0, \boldsymbol{x}_1\right) = \int_0^T dt\, D_t\left(\boldsymbol{x}' \leftarrow \boldsymbol{x}; a_0, \boldsymbol{x}_1\right). \tag{S5.13}$$

By using the expected number of the cells that divide at time $t$ with the division time $a$ and the state $\boldsymbol{x}$, $D_t\left(a, \boldsymbol{x}_1; a_0, \boldsymbol{x}_0\right)$, and $\mathbb{T}_{\mathrm{F}}$, we have

$$D_t\left(\boldsymbol{x}' \leftarrow \boldsymbol{x}; a_0, \boldsymbol{x}_1\right) = 2\mathbb{T}_{\mathrm{F}}(\boldsymbol{x}'|\boldsymbol{x}) \int_0^\infty da\, D_t\left(a, \boldsymbol{x}; a_0, \boldsymbol{x}_0\right), \tag{S5.14}$$

For the rigorous derivation of this equation, see Lemma 1 in the next section. Hence, we have

$$\mathring{N}_T\left(\boldsymbol{x}' \leftarrow \boldsymbol{x}; a_0, \boldsymbol{x}_1\right) = 2\mathbb{T}_{\mathrm{F}}(\boldsymbol{x}'|\boldsymbol{x}) \int_0^T dt \int_0^\infty da\, D_t\left(a, \boldsymbol{x}; a_0, \boldsymbol{x}_0\right). \tag{S5.15}$$

Substituting Eq. (S5.15) into Eq. (S5.12), we obtain

$$\mathbb{T}^{\mathcal{T}_{\boldsymbol{x}}}_{\mathrm{emp}}(\boldsymbol{x}'|\boldsymbol{x}) = \mathbb{T}_{\mathrm{F}}(\boldsymbol{x}'|\boldsymbol{x}). \tag{S5.16}$$

Here, we note the integration part in Eq. (S5.15) is independent of $x'$ and

$$\mathring{N}^{\mathrm{tot}}_T(\boldsymbol{x}) = \int_0^T dt \int_0^\infty da\, D_t\left(a, \boldsymbol{x}; a_0, \boldsymbol{x}_0\right). \tag{S5.17}$$

### S6 A Rigorous proof of the convergence of the empirical distributions

In this section, we present a rigorous proof of the convergence of the empirical distributions for tree sampling. For preparation, we introduce generalizations of Eq. (S4.8), which are called many-to-one formulae [15, 16]. The formulae enable us to calculate the expectation of a summation over a linage tree (many) by the expectation of the retrospective process (one). We first show Eqs. (S5.5) and (S5.14).

**Lemma 1** (many-to-one formula for non-leaf cells)**.** The following equalities hold:

$$D_t\left(\tau, x; a_0, \boldsymbol{x}_1\right) = r_F(\tau, \boldsymbol{x}) N_t\left(\tau, \boldsymbol{x}; a_0, \boldsymbol{x}_1\right), \tag{S6.1}$$

and

$$D_t\left(\boldsymbol{x}' \leftarrow \boldsymbol{x}; a_0, \boldsymbol{x}_1\right) = 2\mathbb{T}_{\mathrm{F}}(\boldsymbol{x}'|\boldsymbol{x}) \int_0^\infty da\, D_t\left(a, \boldsymbol{x}; a_0, \boldsymbol{x}_1\right). \tag{S6.2}$$

*Proof.* In the following, we denote an expected value in capitalized roman and a random variable in calligraphy. We first show Eq. (S6.1). Let a random variable $\mathcal{N}_t\left(a, \boldsymbol{x}_1; a_0, \boldsymbol{x}_0\right)$ be the number of the cells at time $t$ with the age $a$ and the state $\boldsymbol{x}_1$, and $\mathcal{D}_t\left(\tau, \boldsymbol{x}_1; a_0, \boldsymbol{x}_0\right)$ be the number of the non-leaf cells that divides at time $t$ with the division time $\tau$ and the state $\boldsymbol{x}_1$. The notation $i \in \mathcal{N}_t\left(a, \boldsymbol{x}_1; a_0, \boldsymbol{x}_0\right)$ means that the cell $i$ is alive at time $t$ with the age $a$ and the state $\boldsymbol{x}_1$. Let $\mathrm{divides}_i(\tau) := \delta(\tau_i - \tau)$, where the random variable $\tau_i$ is the division time of the cell $i$. By using the following fact about the conditional expectation, $\mathbb{E}[\cdot] = \mathbb{E}\left[\mathbb{E}[\cdot \mid A]\right]$, we obtain

$$D_t\left(\tau, \boldsymbol{x}; a_0, \boldsymbol{x}_1\right) = \mathbb{E}\left[\mathcal{D}_t\left(\tau, \boldsymbol{x}; a_0, \boldsymbol{x}_1\right)\right] = \mathbb{E}\left[\mathbb{E}\left[\sum_{i \in \mathcal{N}_t\left(\tau, \boldsymbol{x}; a_0, \boldsymbol{x}_0\right)} \mathrm{divides}_i(\tau) \middle| \mathcal{N}_t\left(a, \boldsymbol{x}; a_0, \boldsymbol{x}_1\right)\right]\right] \tag{S6.3}$$

$$= \mathbb{E}\left[\sum_{i \in \mathcal{N}_t\left(\tau, \boldsymbol{x}; a_0, \boldsymbol{x}_0\right)} \mathbb{E}\left[\mathrm{divides}_i(\tau) | \mathcal{N}_t\left(a, \boldsymbol{x}; a_0, \boldsymbol{x}_1\right)\right]\right]. \tag{S6.4}$$

Since $\mathrm{divides}_i(\tau)$ and $\mathcal{N}_t\left(a, \boldsymbol{x}; a_0, \boldsymbol{x}_1\right)$ are independent, the following equation holds.

$$\mathbb{E}\left[\mathrm{divides}_i(\tau) | \mathcal{N}_T\left(a, \boldsymbol{x}; a_0, \boldsymbol{x}_1\right)\right] = \mathbb{E}[\mathrm{divides}_i(\tau)] = r(\tau|\boldsymbol{x}), \tag{S6.5}$$

holds. Using this equation in (S6.4), we can obtain the first equality. The second equality can be shown by a similar argument. □

Informally, this proof can be rephrased as follows. Let $\mathcal{N} = \mathcal{N}_t\left(a, \boldsymbol{x}; a_0, \boldsymbol{x}_1\right)$ and $\mathcal{T}$ be a realization of a lineage tree. Then, we get the property of the conditional probability:

$$\mathbb{P}[\mathcal{T}] = \sum_{\mathcal{N}} \mathbb{P}[\mathcal{T}|\mathcal{N}]\mathbb{P}[\mathcal{N}]. \tag{S6.6}$$

Hence, $D_t\left(\tau, \boldsymbol{x}; a_0, \boldsymbol{x}_1\right)$ can be represented as

$$\begin{aligned} D_t\left(\tau, \boldsymbol{x}; a_0, \boldsymbol{x}_1\right) &= \sum_{\mathcal{T}} \sum_{\mathcal{N}} \mathcal{D}_t\left(\tau, \boldsymbol{x}; a_0, x_1\right) \mathbb{P}[\mathcal{T}|\mathcal{N}]\mathbb{P}[\mathcal{N}] \\ &= \sum_{\mathcal{N}} \left[\sum_{\mathcal{T}} \mathcal{D}_t\left(\tau, \boldsymbol{x}; a_0, \boldsymbol{x}_1\right) \mathbb{P}[\mathcal{T}|\mathcal{N}]\right] \mathbb{P}[\mathcal{N}] \\ &= \sum_{\mathcal{N}} \left[\sum_{\mathcal{T}} r(\tau|\boldsymbol{x}) \mathcal{N}_t\left(\tau, \boldsymbol{x}; a_0, \boldsymbol{x}_1\right)\right] \mathbb{P}[\mathcal{N}] \\ &= r(\tau|\boldsymbol{x}) N_t\left[\tau, \boldsymbol{x}; a_0, \boldsymbol{x}_1\right]. \end{aligned} \tag{S6.7}$$

We next show variants of Lemma 1. Let $\mathring{N}^c_T\left((\tau', \boldsymbol{x}') : (\tau'', \boldsymbol{x}''); a_0, \boldsymbol{x}_1\right)$ be the expected number of the pairs $(i, j)$ of the non-leaf cells satisfying:

$$\begin{aligned} &(1) \text{ the cell } i \text{ is an ancestor of the cell } j; \\ &(2) \text{ the cell } i \text{ has division time } \tau' \text{ and the state } \boldsymbol{x}'; \\ &(3) \text{ the cell } j \text{ does the division time } \tau'' \text{ and the state } \boldsymbol{x}''. \end{aligned} \tag{S6.8}$$

Similarly, let $\mathring{N}^u_T\left((\tau', \boldsymbol{x}') : (\tau'', \boldsymbol{x}''); a_0, \boldsymbol{x}_1\right)$ be the expected number of the pairs of the non-leaf cell satisfying:

$$\begin{aligned} &(1) \text{ the cell } i \text{ is neither an ancestor nor a descendant of the cell } j; \\ &(2) \text{ the cell } i \text{ has the division time } \tau' \text{ and the state } \boldsymbol{x}'; \\ &(3) \text{ the cell } j \text{ does the division time } \tau'' \text{ and the state } \boldsymbol{x}''. \end{aligned} \tag{S6.9}$$

**Lemma 2** (many-to-one formula for folks)**.** The following equality holds:

$$\begin{aligned} &\mathring{N}^c_T\left((\tau', \boldsymbol{x}') : (\tau'', \boldsymbol{x}''); a_0, \boldsymbol{x}_1\right) \\ &= 2\int_0^T dt\, D_t\left(\tau', \boldsymbol{x}'; a_0, \boldsymbol{x}_1\right) \sum_{\boldsymbol{x}''' \in \Omega} \mathring{N}_{T-t}\left(\tau'', \boldsymbol{x}''; 0, \boldsymbol{x}'''\right) \mathbb{T}_{\mathrm{F}}(\boldsymbol{x}'''|\boldsymbol{x}'), \end{aligned} \tag{S6.10}$$

and

$$
\begin{aligned}
&\mathring{N}^{u}_{T}\left((\tau', \boldsymbol{x}') : (\tau'', \boldsymbol{x}''); a_0, \boldsymbol{x}_1\right) \\
&= 2\int_0^T dt \int_0^\infty d\tau''' \sum_{\boldsymbol{x}'''\in\Omega} D_t\left(\tau''', \boldsymbol{x}'''; a_0, \boldsymbol{x}_1\right) \left[\sum_{\boldsymbol{x}''''\in\Omega} \mathring{N}_{T-t}\left(\tau', \boldsymbol{x}'; 0, \boldsymbol{x}''''\right) \mathbb{T}_{\mathrm{F}}(\boldsymbol{x}''''|\boldsymbol{x}''')\right] \\
&\quad \cdot \left[\sum_{\boldsymbol{x}''''\in\Omega} \mathbb{T}_{\mathrm{F}}(\boldsymbol{x}''''|\boldsymbol{x}''')\mathring{N}_{T-t}\left(\tau'', \boldsymbol{x}''; 0, \boldsymbol{x}''''\right) \mathbb{T}_{\mathrm{F}}(\boldsymbol{x}''''|\boldsymbol{x}''')\right]. \qquad \text{(S6.11)}
\end{aligned}
$$

Before the proof, we explain the intuitive meaning of the equations. We first explain Eq. (S6.10). We start with evaluating the expected number of the cell $i$ and the cell $j$ satisfying (i) the conditions (S6.8) and (ii) the cell $i$ divides at $t$. The factor $D_t\left(\tau', \boldsymbol{x}'; a_0, \boldsymbol{x}_1\right)$ is the expected number of such cell $i$. Then, $2\ \mathbb{T}_{\mathrm{F}}(\boldsymbol{x}'''|\boldsymbol{x}')$ is the expected number of the daughter cells of the cell $i$ with state $\boldsymbol{x}'''$. Hence, $2\mathring{N}_{T-t}\left(\tau'', \boldsymbol{x}''; 0, \boldsymbol{x}'''\right) \mathbb{T}_{\mathrm{F}}(\boldsymbol{x}'''|\boldsymbol{x}')$ is the expected number of such cell $j$. All in all, we know that such expected number is $2D_t\left(\tau', \boldsymbol{x}'; a_0, \boldsymbol{x}_1\right) \sum_{\boldsymbol{x}'''\in\Omega} \mathring{N}_{T-t}\left(\tau'', \boldsymbol{x}''; 0, \boldsymbol{x}'''\right) \mathbb{T}_{\mathrm{F}}(\boldsymbol{x}'''|\boldsymbol{x}')$. Summing up this expected number by the integration $\int_0^T dt$, we have the total expected number of the cell $i$ and the cell $j$ satisfying the conditions (S6.8).

We can give a similar interpretation of Eq. (S6.11). We start with evaluating the expected number of the cell $i$ and the cell $j$ satisfying (i) the conditions (S6.9) and (ii) the latest common ancestor of the cell $i$ and the cell $j$, which we call cell $w$, divides at $t$ with the age $\tau'''$ and the state $\boldsymbol{x}'''$. The factor $D_t\left(\tau''', \boldsymbol{x}'''; a_0, \boldsymbol{x}_1\right)$ is the expected number of such cell $w$. The factor $\sum_{\boldsymbol{x}''''\in\Omega} \mathring{N}_{T-t}\left(\tau', \boldsymbol{x}'; 0, \boldsymbol{x}''''\right) \mathbb{T}_{\mathrm{F}}(\boldsymbol{x}''''|\boldsymbol{x}''')$ and $\sum_{\boldsymbol{x}''''\in\Omega} \mathring{N}_{T-t}\left(\tau'', \boldsymbol{x}''; 0, \boldsymbol{x}''''\right) \mathbb{T}_{\mathrm{F}}(\boldsymbol{x}''''|\boldsymbol{x}''')$ is the expected number of such cell $i$ and cell $j$, respectively. The prefactor 2 stands for the arbitrariness for assigning the label $i$ and the label $j$ to the descendants of the two daughters of the cell $w$. Hence, summing up by $\int_0^T dt \int_0^\infty d\tau'''$, we have the total expected number.

*Proof.* We first show Eq. (S6.10). The random variable $\mathring{\mathcal{N}}^{c}_{T}\left((\tau', \boldsymbol{x}') : (\tau'', \boldsymbol{x}''); a_0, \boldsymbol{x}_1\right)$ is the number of the pairs $(i, j)$ of the non-leaf cells satisfying the conditions (S6.8). Then, we have

$$
\mathring{N}^{c}_{T}\left((\tau', \boldsymbol{x}') : (\tau'', \boldsymbol{x}''); a_0, \boldsymbol{x}_1\right) = \mathbb{E}\left[\mathring{\mathcal{N}}^{c}_{T}\left((\tau', \boldsymbol{x}') : (\tau'', \boldsymbol{x}''); a_0, \boldsymbol{x}_1\right)\right]. \qquad \text{(S6.12)}
$$

By focasing on the time when the ancestor cell divides, we have

$$
\mathring{\mathcal{N}}^{c}_{T}\left((\tau', \boldsymbol{x}') : (\tau'', \boldsymbol{x}''); a_0, \boldsymbol{x}_1\right) = \int_0^T dt \sum_{i\in\mathcal{D}_t(\tau', \boldsymbol{x}'; a_0, \boldsymbol{x}_1)} \mathcal{N}_1(i), \qquad \text{(S6.13)}
$$

where $\mathcal{N}_1(i)$ is the number of the descendants of a cell $i \in \mathcal{D}_t\left(\tau', \boldsymbol{x}'; a_0, \boldsymbol{x}_1\right)$ with the division time $\tau''$ and the state $\boldsymbol{x}''$. By substituting this equality into Eq. (S6.12), we obtain

$$
\begin{aligned}
\mathring{N}^{c}_{T}\left((\tau', \boldsymbol{x}') : (\tau'', \boldsymbol{x}''); a_0, \boldsymbol{x}_1\right) &= \mathbb{E}\left[\int_0^T dt \sum_{i\in\mathcal{D}_t(\tau', \boldsymbol{x}'; a_0, \boldsymbol{x}_1)} \mathcal{N}_1(i)\right], \\
&= \mathbb{E}\left[\int_0^T dt \sum_{i\in\mathcal{D}_t(\tau', \boldsymbol{x}'; a_0, \boldsymbol{x}_1)} \mathbb{E}\left[\mathcal{N}_1(i) | i \in \mathcal{D}_t\left(\tau', \boldsymbol{x}'; a_0, \boldsymbol{x}_1\right)\right]\right].
\end{aligned} \qquad \text{(S6.14)}
$$

We evaluate $\mathbb{E}\left[\mathcal{N}_1(i) | i \in \mathcal{D}_t\left(\tau', \boldsymbol{x}'; a_0, \boldsymbol{x}_1\right)\right]$. Since the cell $i$ has two daughters and each has state $\boldsymbol{x}'''$ with the probability $\mathbb{T}_{\mathrm{F}}(\boldsymbol{x}'''|\boldsymbol{x}')$, we obtain

$$
\mathbb{E}\left[\mathcal{N}_1(i) | i \in \mathcal{D}_t\left(\tau', \boldsymbol{x}'; a_0, \boldsymbol{x}_1\right)\right] = 2\sum_{\boldsymbol{x}'''\in\Omega} \mathring{N}_{T-t}\left(\tau'', \boldsymbol{x}''; 0, \boldsymbol{x}'''\right) \mathbb{T}_{\mathrm{F}}(\boldsymbol{x}'''|\boldsymbol{x}'). \qquad \text{(S6.15)}
$$

Notice that this value is dependent not on $i$ but on $t$. Hence, we have

$$
\begin{aligned}
&\mathring{N}^{c}_{T}\left((\tau', \boldsymbol{x}') : (\tau'', \boldsymbol{x}''); a_0, \boldsymbol{x}_1\right) \\
&= \mathbb{E}\left[\int_0^T dt \sum_{i \in \mathcal{D}_t(\tau', \boldsymbol{x}'; a_0, \boldsymbol{x}_1)} 2 \sum_{\boldsymbol{x}''' \in \Omega} \mathring{N}_{T-t}\left(\tau'', \boldsymbol{x}''; 0, \boldsymbol{x}'''\right) \mathbb{T}_{\mathrm{F}}(\boldsymbol{x}'''|\boldsymbol{x}')\right] \\
&= 2\int_0^T dt \sum_{\boldsymbol{x}''' \in \Omega} \mathring{N}_{T-t}\left(\tau'', \boldsymbol{x}''; 0, \boldsymbol{x}'''\right) \mathbb{T}_{\mathrm{F}}(\boldsymbol{x}'''|\boldsymbol{x}') \mathbb{E}\left[\mathcal{D}_t\left(\tau', \boldsymbol{x}'; a_0, \boldsymbol{x}_1\right)\right] \\
&= 2\int_0^T dt\, D_t\left(\tau', \boldsymbol{x}'; a_0, \boldsymbol{x}_1\right) \sum_{x''' \in \Omega} \mathring{N}_{T-t}\left(\tau'', \boldsymbol{x}''; 0, \boldsymbol{x}'''\right) \mathbb{T}_{\mathrm{F}}(\boldsymbol{x}'''|\boldsymbol{x}'). \qquad \text{(S6.16)}
\end{aligned}
$$

We next prove Eq. (S6.11). The random variable $\mathring{\mathcal{N}}^{u}_{T}\left((\tau', \boldsymbol{x}') : (\tau'', \boldsymbol{x}''); a, \boldsymbol{x}\right)$ is the number of the pairs $(i, j)$ satisfying the conditions (S6.9). Then, we have

$$
\mathring{N}^{u}_{T}\left((\tau', \boldsymbol{x}') : (\tau'', \boldsymbol{x}''); a_0, \boldsymbol{x}_1\right) = \mathbb{E}\left[\mathring{\mathcal{N}}^{u}_{T}\left((\tau', \boldsymbol{x}') : (\tau'', \boldsymbol{x}''); a_0, \boldsymbol{x}_1\right)\right]. \qquad \text{(S6.17)}
$$

By focusing on the time when the latest common ancestor divides and its state, we have

$$
\mathring{\mathcal{N}}^{u}_{T}\left((\tau', \boldsymbol{x}') : (\tau'', \boldsymbol{x}''); a_0, \boldsymbol{x}_1\right) = \int_0^T dt \sum_{w \in \mathcal{D}_t(\tau''', \boldsymbol{x}''; a_0, \boldsymbol{x}_1)} \mathcal{N}_2(w), \qquad \text{(S6.18)}
$$

where $\mathcal{N}_2(w)$ is the number of the pairs $(i, j)$ of the descendants of cell $w \in \mathcal{D}_t\left(\tau''', \boldsymbol{x}'''; a_0, \boldsymbol{x}_1\right)$ satisfying: (1) the cell $i$ and the cell $j$ are descendants of the different daughters of the cell $w$. (2) the cell $i$ has the division time $\tau'$ and the state $\boldsymbol{x}'$; (3) the cell $j$ has the division time $\tau''$ and the state $\boldsymbol{x}''$. Then, we obtain

$$
\mathring{N}^{u}_{T}\left((\tau', \boldsymbol{x}') : (\tau'', \boldsymbol{x}''); a_0, \boldsymbol{x}_1\right) \qquad \text{(S6.19)}
$$

$$
= \mathbb{E}\left[\int_0^T dt \int_0^\infty d\tau''' \sum_{\boldsymbol{x}''' \in \Omega} \sum_{w \in \mathcal{D}_t(\tau''', \boldsymbol{x}'''; a_0, \boldsymbol{x}_1)} \mathbb{E}\left[\mathcal{N}_2(w) | w \in \mathcal{D}_t\left(\tau''', \boldsymbol{x}'''; a_0, \boldsymbol{x}_1\right)\right]\right]. \qquad \text{(S6.20)}
$$

We evaluate $\mathbb{E}\left[\mathcal{N}_2(w) | w \in \mathcal{D}_t\left(\tau''', \boldsymbol{x}'''; a_0, \boldsymbol{x}_1\right)\right]$ . Since the two sub-lineages that consist of the descendants of the two daughters are independent of the cell $w$, we have

$$
\mathcal{N}_2(w) = 2\left(\sum_{\boldsymbol{x}'''' \in \Omega} \mathbb{T}_{\mathrm{F}}(\boldsymbol{x}''''|\boldsymbol{x}''') \mathring{N}_{T-t}\left(\tau', \boldsymbol{x}'; 0, \boldsymbol{x}''''\right)\right)\left(\sum_{\boldsymbol{x}'''' \in \Omega} \mathbb{T}_{\mathrm{F}}(\boldsymbol{x}''''|\boldsymbol{x}''') \mathring{N}_{T-t}\left(\tau'', \boldsymbol{x}''; 0, \boldsymbol{x}''''\right)\right). \qquad \text{(S6.21)}
$$

Hence we obtain

$$
\begin{aligned}
\mathring{N}^{u}_{T}\left((\tau', \boldsymbol{x}') : (\tau'', \boldsymbol{x}''); a_0, \boldsymbol{x}_1\right) = 2\int_0^T dt \int_0^\infty d\tau''' \sum_{\boldsymbol{x}''' \in \Omega} D_t\left(\tau''', \boldsymbol{x}'''; a_0, \boldsymbol{x}_1\right) \\
\left[\sum_{\boldsymbol{x}'''' \in \Omega} \mathbb{T}_{\mathrm{F}}(\boldsymbol{x}''''|\boldsymbol{x}''') \mathring{N}_{T-t}\left(\tau', \boldsymbol{x}'; 0, \boldsymbol{x}''''\right)\right] \\
\left[\sum_{\boldsymbol{x}'''' \in \Omega} \mathbb{T}_{\mathrm{F}}(\boldsymbol{x}''''|\boldsymbol{x}''') \mathring{N}_{T-t}\left(\tau'', \boldsymbol{x}''; 0, \boldsymbol{x}''''\right)\right]. \qquad \text{(S6.22)}
\end{aligned}
$$

□

We need three assumptions to prove the convergence of $\pi_{\mathrm{emp}}$ and $\mathbb{T}_{\mathrm{emp}}$. The first assumption is that the growth rate must satisfy

$$
\log|\mathcal{T}_{\boldsymbol{x}}| - \lambda T = o(T), \qquad \text{(S6.23)}
$$

almost surely. The second assumption is the geometric ergodicity of the retrospective process. Recall that $p^B_{\rm st}(a,\boldsymbol{x})$ is the stationary measure of the retrospective process. We assume that

$$\left|\int da \sum_{\boldsymbol{x}\in\Omega} f(a,\boldsymbol{x})p^B_t(a,\boldsymbol{x};a_0,\boldsymbol{x}_1) - \int da \sum_{\boldsymbol{x}\in\Omega} f(a,\boldsymbol{x})p^B_{\rm st}(a,\boldsymbol{x})\right| \leq \text{const.} \times \|f\|_\infty e^{-\rho t}, \tag{S6.24}$$

for all bounded function $f$ and all initial condition $(a_0,\boldsymbol{x}_1)\in\mathbb{R}\times\Omega$. In the following, the notation const. means a constant factor with respect to time variables other than $a_0$. The third assumption is that the rate $r_B(a\mid\boldsymbol{x})$ is bounded. These assumptions must be checked depending on the details of the model. In practice, the assumption about the growth rate can be checked experimentally. The sufficient condition for the second assumptions is given in Ref. [19]. We mention that the second property is intuitive. Markov processes with finite state spaces satisfy the geometric ergodicity if the transition matrix is ergodic owing to the Peron-Frobenius theorem. Considering the analogy of this fact, we can naturally expect that the geometric ergodicity holds if $\mathbb{T}_{\rm F}$ is ergodic and $\pi_{\rm F}$ is a non-singular distribution. It is easy to see that the third assumption is satisfied if $\pi_{\rm F}$ is a log-normal distribution or a gamma distribution with shape parameter not less than 1.

In the proof of the convergence of the empirical distributions, we only need the following coarse inequalities obtained by neglecting constant factors in the many-to-one formulae.

**Lemma 3.** Let $\boldsymbol{x}\in\Omega$ and $f\colon\mathbb{R}\to\mathbb{R}$ be a bounded function with $\int_0^\infty da\, f(a)r_B(a\mid\boldsymbol{x})p^B_{\rm st}(a,\boldsymbol{x})=0$. Then, the following inequalities hold under the three assumptions above;

$$\left|\int_0^\infty d\tau''\, f(\tau'')\mathring{N}^c_T\left((\tau',\boldsymbol{x}):(\tau'',\boldsymbol{x});a_0,\boldsymbol{x}_1\right)\right| \leq \text{const.}\times\int_0^T dt\, e^{\lambda t}\int_0^{T-t} ds\, e^{(\lambda-\rho)s}, \tag{S6.25}$$

and

$$\left|\int_0^\infty d\tau'\int_0^\infty d\tau''\, f(\tau')f(\tau'')\mathring{N}^u_T\left((\tau',\boldsymbol{x}):(\tau'',\boldsymbol{x});a_0,\boldsymbol{x}_0\right)\right| \tag{S6.26}$$

$$\leq \text{const.}\times\int_0^T dt\, e^{\lambda t}\left[\int_0^{T-t} ds \sum_{\boldsymbol{x}'''\in\Omega} e^{(\lambda-\rho)s}\right]^2. \tag{S6.27}$$

*Proof.* We first prove Eq. (S6.25). By neglecting constant factors and using Eqs. (S5.6), (S6.1), and (S6.10), we have

$$\mathring{N}^c_T\left((\tau',\boldsymbol{x}'):(\tau'',\boldsymbol{x}'');a_0,\boldsymbol{x}_1\right) = \text{const.}\times\int_0^T dt\, e^{\lambda t}\sum_{\boldsymbol{x}'''\in\Omega}\int_0^{T-t} ds\int_0^\infty d\tau''\, e^{\lambda s}r_B(\tau''\mid\boldsymbol{x}'')p^B_s(\tau'',\boldsymbol{x}'';0,\boldsymbol{x}'''). \tag{S6.28}$$

By integrating it with respect to $f$ and by setting $\boldsymbol{x}'=\boldsymbol{x}''=\boldsymbol{x}$, we have

$$\left|\int_0^\infty d\tau''\, f(\tau'')\mathring{N}^c_T\left((\tau',\boldsymbol{x}):(\tau'',\boldsymbol{x});a_0,\boldsymbol{x}_1\right)\right| \tag{S6.29}$$

$$\leq \text{const.}\times\int_0^T dt\, e^{\lambda t}\sum_{\boldsymbol{x}'''\in\Omega}\int_0^{T-t} ds\, f(\tau'')e^{\lambda s}r_B(\tau''\mid\boldsymbol{x}'')p^B_s(\tau'',\boldsymbol{x}'';0,\boldsymbol{x}'''). \tag{S6.30}$$

By the assumption (S6.24), we obtain in sufficiently large $T$

$$\left|\int_0^\infty d\tau''\, f(\tau'')\mathring{N}^c_T\left((\tau',\boldsymbol{x}):(\tau'',\boldsymbol{x});a_0,\boldsymbol{x}_1\right)\right| \tag{S6.31}$$

$$\leq \text{const.}\times\left|\int_0^T dt\, e^{\lambda t}\int_0^{T-t} ds\int_0^\infty d\tau''\, f(\tau'')e^{\lambda s}r_B(\tau''\mid\boldsymbol{x}'')p^B_{\rm st}(\tau'',\boldsymbol{x})\right| \tag{S6.32}$$

$$+\,\text{const.}\times\left|\int_0^T dt\, e^{\lambda t}\int_0^{T-t} ds\int_0^\infty d\tau''\, e^{-\rho s}\right| \tag{S6.33}$$

$$\leq \text{const.}\times\int_0^T dt\, e^{\lambda t}\int_0^{T-t} ds\, e^{(\lambda-\rho)s}. \tag{S6.34}$$

Here, we used $p_s^B(\tau'', \boldsymbol{x}''; 0, \boldsymbol{x}''') = \sum_{\boldsymbol{x}} \delta_{\boldsymbol{x},\boldsymbol{x}''} p_s^B(\tau'', \boldsymbol{x}; 0, \boldsymbol{x}''')$.

We next prove Eq. (S6.26). By neglecting constant factors and using Eqs. (S5.6), (S6.1), and (S6.11), we obtain

$$\begin{aligned} &\mathring{N}_T^u((\tau', \boldsymbol{x}) : (\tau'', \boldsymbol{x}); a_0, \boldsymbol{x}_0) \\ &= \text{const.} \times \int_0^T dt\, e^{\lambda t} \left[ \int_0^{T-t} ds \sum_{\boldsymbol{x}''' \in \Omega} e^{\lambda s} r_B(\tau' \mid \boldsymbol{x}') p_s^B(\tau', \boldsymbol{x}'; 0, \boldsymbol{x}''') \right] \\ &\qquad \left[ \int_0^{T-t} ds \sum_{\boldsymbol{x}''' \in \Omega} e^{\lambda s} r_B(\tau'' \mid \boldsymbol{x}'') p_s^B(\tau'', \boldsymbol{x}''; 0, \boldsymbol{x}''') \right]. \end{aligned} \tag{S6.35}$$

By integrating it with respect to $f$, we have

$$\begin{aligned} &\left| \int_0^\infty d\tau' \int_0^\infty d\tau''\, f(\tau') f(\tau'') \mathring{N}_T^u((\tau', \boldsymbol{x}) : (\tau'', \boldsymbol{x}); a_0, \boldsymbol{x}_0) \right| \\ &\leq \text{const.} \times \int_0^T dt\, e^{\lambda t} \left[ \int_0^{T-t} ds \sum_{\boldsymbol{x}''' \in \Omega} e^{(\lambda - \rho)s} \right]^2. \end{aligned} \tag{S6.36}$$

□

We are ready to prove the convergence of the empirical distributions.

**Theorem 4.** Under the three assumptions above, the empirical distributions converges in distribution:

$$\pi_{\text{emp}}^{\mathcal{T}_{\boldsymbol{x}}}(\tau|\boldsymbol{x}) \to \pi_{\text{B}}(\tau|\boldsymbol{x}), \tag{S6.37}$$

$$\mathbb{T}_{\text{emp}}^{\mathcal{T}_{\boldsymbol{x}}}(\boldsymbol{x}'|\boldsymbol{x}) \to \mathbb{T}_{\text{F}}(\boldsymbol{x}'|\boldsymbol{x}). \tag{S6.38}$$

Rigorously, Eq. (S6.37) means

$$\frac{1}{|\mathcal{T}_{\boldsymbol{x}}|} \sum_{i \in \mathcal{T}_{\boldsymbol{x}}} f(\tau_i) \to \int_0^\infty d\tau\, f(\tau) \pi_{\text{B}}(\tau|\boldsymbol{x}), \tag{S6.39}$$

for all $\tau$ and bounded function $f$.

*Proof.* For simplicity, we just denote $\pi_{\text{emp}}^{\mathcal{T}_{\boldsymbol{x}}}(\tau|\boldsymbol{x})$ by $\pi_{\text{emp}}(\tau|\boldsymbol{x})$ and $\mathbb{T}_{\text{emp}}^{\mathcal{T}_{\boldsymbol{x}}}(\boldsymbol{x}'|\boldsymbol{x})$ by $\mathbb{T}_{\text{emp}}(\boldsymbol{x}'|\boldsymbol{x})$. We first prove Eq. (S6.37). It suffices to show this convergence for all bounded function $f$ satisfying $\int_0^\infty da\, f(a) p_{\text{st}}^B(a, \boldsymbol{x}) = 0$, because (i) Eq. (S6.39) is linear in $f$, (ii) $f_0(\tau) = f(\tau) - \int_0^\infty da\, f(a) r_B(a \mid \boldsymbol{x}) p_{\text{st}}^B(a, \boldsymbol{x})$ is a bounded function satisfying $\int_0^\infty da\, f_0(a) r_B(a \mid \boldsymbol{x}) p_{\text{st}}^B(a, \boldsymbol{x}) = 0$. This assumption implies that $\int_0^\infty da\, f(a) \pi_{\text{B}}(a|\boldsymbol{x}) = 0$ since $r_B(a \mid \boldsymbol{x}) p_{\text{st}}^B(a, \boldsymbol{x})$ is the distribution of division time of cells with state $\boldsymbol{x}$, which is also characterized by $\pi_{\text{B}}(a|\boldsymbol{x})$. Hence, it suffices to show that

$$\mathbb{E}\left[ \left( \frac{1}{|\mathcal{T}_{\boldsymbol{x}}|} \sum_{i \in \mathcal{T}_{\boldsymbol{x}}} f(\tau_i) \right)^2 \right] \to 0. \tag{S6.40}$$

Let $\mathcal{C}$ be the set of the pairs $(i, j)$ of the non-leaf cells, where the cell $i$ is the ancestor of the cell $j$, and $\mathcal{U}$ be the set of the pairs $(i, j)$, where the cell $i$ is neither the ancestor nor the descendant of the cell $j$. From Hölder's inequality,

$$\mathbb{E}\left[ \left( \frac{1}{|\mathcal{T}_{\boldsymbol{x}}|} \sum_{i \in \mathcal{T}_{\boldsymbol{x}}} f(\tau_i) \right)^2 \right] \leq \frac{e^{2\lambda T}}{|\mathcal{T}_{\boldsymbol{x}}|^2} \cdot e^{-2\lambda T} \left[ (I) + (II) + (III) \right], \tag{S6.41}$$

holds, where

$$(I) = \mathbb{E}\left[\sum_{i\in\mathcal{T}_{\boldsymbol{x}}} f(\tau_i)^2\right], \tag{S6.42}$$

$$(II) = 2\mathbb{E}\left[\sum_{(i,j)\in\mathcal{C}} f(\tau_i)f(\tau_j)\right], \tag{S6.43}$$

$$(III) = \mathbb{E}\left[\sum_{(i,j)\in\mathcal{U}} f(\tau_i)f(\tau_j)\right]. \tag{S6.44}$$

$$\tag{S6.45}$$

Notice that the factor $e^{2\lambda T}/|\mathcal{T}_{\boldsymbol{x}}|^2$ is $e^{o(T)}$ from the assumption of the growth rate. Hence, it suffices to show that $2\lambda T - \log[(I)]$, $2\lambda T - \log[(II)]$, and $2\lambda T - \log[(III)]$ are positive at sufficiently large $T$ and $\Omega(T)$. Here, a positive function $h(T)$ is $\Omega(T)$ when $\lim_{T\to\infty} T/h(T)$ converges. Since

$$(I) = \int_0^\infty d\tau\, f^2(\tau)\mathring{N}_T\,(\tau,\boldsymbol{x};a_0,\boldsymbol{x}_1) \leq \|f\|_\infty^2|\mathcal{T}|, \tag{S6.46}$$

we obtain $\log[(I)] = \lambda T + o(T)$. Hence, $2\lambda T - \log[(I)]$ is positive at sufficiently large $T$ and $\Omega(T)$. For the second term, Eq. (S6.25) implies

$$\begin{aligned}|(II)| &= 2\left|\int d\tau' \int d\tau'' f(\tau')f(\tau'')\mathring{N}_T^c\,((\tau',\boldsymbol{x}):(\tau'',\boldsymbol{x});a_0,\boldsymbol{x}_0)\right| \\ &\leq \text{const.}\times \int_0^T dt e^{\lambda t}\int_0^{T-t} ds\, e^{(\lambda-\rho)s}.\end{aligned} \tag{S6.47}$$

By a direct calculation, we can show that $2\lambda T - \log[(II)] = \Omega(T)$. Hence, $2\lambda T - \log[(II)]$ is positive at sufficiently large $T$ and $\Omega(T)$. For the third term, Eq. (S6.26) implies

$$\begin{aligned}|(III)| &= 2\left|\int d\tau' \int d\tau''\, f(\tau')f(\tau'')\mathring{N}_T^u\,((\tau',\boldsymbol{x}):(\tau'',\boldsymbol{x});a_0,\boldsymbol{x}_0)\right| \\ &\leq \text{const.}\times \left|\int_0^T dt\, e^{\lambda t}\left(\int_0^{T-t} ds\, e^{(\lambda-\rho)}\right)^2\right|.\end{aligned} \tag{S6.48}$$

By direct calculation, we have $2\lambda T - \log[(III)] = \Omega(T)$. Hence, $2\lambda T - \log[(III)]$ is positive at sufficiently large $T$ and $\Omega(T)$. These completes the proof of Eq. (S6.37).

We next prove Eq. (S6.38) by a similar argument. For notational simplicity, we denote the Kronecker's delta by $\delta(\cdot,\cdot)$ in the following. Let $g_{\boldsymbol{x}'\leftarrow\boldsymbol{x}}(\boldsymbol{x}_l,\boldsymbol{x}_r) := 1/2(\delta(\boldsymbol{x}_l,\boldsymbol{x}')+\delta(\boldsymbol{x}_r,\boldsymbol{x}')) - \mathbb{T}_{\mathrm{F}}(\boldsymbol{x}'|\boldsymbol{x})$. Notice that $\|g\|_\infty \leq 1$. Eq. (S6.38) is equivalent to the following condition:

$$\frac{1}{|\mathcal{T}_{\boldsymbol{x}}|}\sum_{i\in\mathcal{T}_{\boldsymbol{x}}} g(\boldsymbol{x}_{l(i)},\boldsymbol{x}_{r(i)}) \to 0 \tag{S6.49}$$

where $l(i)$ and $r(i)$ denote the daughters of the cell $i$. Hence, it suffices to show that

$$\mathbb{E}\left[\left[\frac{1}{|\mathcal{T}_{\boldsymbol{x}}|}\sum_{i\in\mathcal{T}_x} g_{\boldsymbol{x}'\leftarrow\boldsymbol{x}}(\boldsymbol{x}_{l(i)},\boldsymbol{x}_{r(i)})\right]^2\right] \to 0. \tag{S6.50}$$

By Hölder's inequality,

$$\mathbb{E}\left[\left[\frac{1}{|\mathcal{T}_{\boldsymbol{x}}|}\sum_{i\in\mathcal{T}_x} g_{\boldsymbol{x}'\leftarrow\boldsymbol{x}}(\boldsymbol{x}_{l(i)},\boldsymbol{x}_{r(i)})\right]^2\right] \leq \frac{e^{2\lambda T}}{|\mathcal{T}_{\boldsymbol{x}}|^2}\cdot e^{-2\lambda T}\,[(I)+(II)+(III)] \tag{S6.51}$$

where

$$(I) = \mathbb{E}\left[\sum_{i\in\mathcal{T}_{\boldsymbol{x}}} g_{\boldsymbol{x}'\leftarrow\boldsymbol{x}}(\boldsymbol{x}_{l(i)}, \boldsymbol{x}_{r(i)})^2\right], \tag{S6.52}$$

$$(II) = 2\mathbb{E}\left[\sum_{(i,j)\in\mathcal{C}} g_{\boldsymbol{x}'\leftarrow\boldsymbol{x}}(\boldsymbol{x}_{l(i)}, \boldsymbol{x}_{r(i)}) g_{\boldsymbol{x}'\leftarrow\boldsymbol{x}}(\boldsymbol{x}_{l(j)}, \boldsymbol{x}_{r(j)})\right], \tag{S6.53}$$

$$(III) = \mathbb{E}\left[\sum_{(i,j)\in\mathcal{U}} g_{\boldsymbol{x}'\leftarrow\boldsymbol{x}}(\boldsymbol{x}_{l(i)}, \boldsymbol{x}_{r(i)}) g_{\boldsymbol{x}'\leftarrow\boldsymbol{x}}(\boldsymbol{x}_{l(j)}, \boldsymbol{x}_{r(j)})\right]. \tag{S6.54}$$

$$\tag{S6.55}$$

Hence, it suffices to show that $e^{-2\lambda T}(I)$, $e^{-2\lambda T}(II)$, and $e^{-2\lambda T}(III)$ converges to 0. We first discuss the first term. Let $\mathring{N}_t\left(\boldsymbol{x}_2, \boldsymbol{x}_3 \leftarrow \tau, \boldsymbol{x}; a_0, \boldsymbol{x}_1\right)$ be the number of the non-leaf cells satisfying: (1) their state and division time are $\boldsymbol{x}$ and $\tau$; (2) the cells divide at time $t$; (3) the state of their daughters are $\boldsymbol{x}_2$ and $\boldsymbol{x}_3$. Then,

$$\mathring{N}_t\left(\boldsymbol{x}_2, \boldsymbol{x}_3 \leftarrow \tau, \boldsymbol{x}; a_0, \boldsymbol{x}_1\right) = \mathbb{T}_{\mathrm{F}}(\boldsymbol{x}_2|\boldsymbol{x})\mathbb{T}_{\mathrm{F}}(\boldsymbol{x}_3|\boldsymbol{x})\mathring{N}_t\left(\tau, \boldsymbol{x}; a_0, \boldsymbol{x}_1\right). \tag{S6.56}$$

since the type switching of the daughters are conditionally independent given the state of their mother (cf. Eq. S6.3). Hence, we can evaluate the first factor as

$$\begin{aligned}
|(I)| &= \left|\int d\tau\, \mathring{N}_t\left(\tau, \boldsymbol{x}; a_0, \boldsymbol{x}_1\right) \sum_{\boldsymbol{x}_{l(i)}, \boldsymbol{x}_{r(i)}\in\Omega} g^2_{\boldsymbol{x}'\leftarrow\boldsymbol{x}}(\boldsymbol{x}_{l(i)}, \boldsymbol{x}_{r(i)})\mathbb{T}_{\mathrm{F}}(\boldsymbol{x}_{l(i)}|\boldsymbol{x})\mathbb{T}_{\mathrm{F}}(\boldsymbol{x}''|\boldsymbol{x})\right| \\
&\le \left|\int d\tau\, \mathring{N}_t\left(\tau, \boldsymbol{x}; a_0, \boldsymbol{x}_1\right)\right| \\
&\le |\mathcal{T}_{\boldsymbol{x}}| \\
&= e^{\lambda T + o(T)}.
\end{aligned} \tag{S6.57}$$

In the last equation, we use the assumption on the growth rate. Hence, $e^{-2\lambda T}(I)$ converges to 0. We next discuss the second term $(II)$. We generalize Eq. (S6.10) by a similar argument and obtain

$$\begin{aligned}
(II) = 2\int_0^T dt \int d\tau &\sum_{\boldsymbol{x}_{l(i)}, \boldsymbol{x}_{r(i)}\in\Omega} D_t\left(\tau, \boldsymbol{x}; a_0, \boldsymbol{x}_1\right) g_{\boldsymbol{x}'\leftarrow\boldsymbol{x}}(\boldsymbol{x}_{l(i)}, \boldsymbol{x}_{r(i)})\mathbb{T}_{\mathrm{F}}(\boldsymbol{x}_{l(i)}|\boldsymbol{x})\mathbb{T}_{\mathrm{F}}(\boldsymbol{x}_{r(i)}|\boldsymbol{x}) \\
\int d\tau' &\left(\mathring{N}_{T-t}\left(\tau', \boldsymbol{x}; 0, \boldsymbol{x}_{l(i)}\right) + \mathring{N}_{T-t}\left(\tau', \boldsymbol{x}; 0, \boldsymbol{x}_{r(i)}\right)\right) \\
&\sum_{\boldsymbol{x}_{l(j)}, \boldsymbol{x}_{r(j)}\in\Omega} g_{\boldsymbol{x}'\leftarrow\boldsymbol{x}}(\boldsymbol{x}_{l(j)}, \boldsymbol{x}_{r(j)})\mathbb{T}_{\mathrm{F}}(\boldsymbol{x}_{l(j)}|\boldsymbol{x})\mathbb{T}_{\mathrm{F}}(\boldsymbol{x}_{r(j)}|\boldsymbol{x}).
\end{aligned} \tag{S6.58}$$

Since the last sum $\sum_{\boldsymbol{x}_{l(j)}, \boldsymbol{x}_{r(j)}\in\Omega} g_{\boldsymbol{x}'\leftarrow\boldsymbol{x}}(\boldsymbol{x}_{l(j)}, \boldsymbol{x}_{r(j)})\mathbb{T}_{\mathrm{F}}(\boldsymbol{x}_{l(j)}|\boldsymbol{x})\mathbb{T}_{\mathrm{F}}(\boldsymbol{x}_{r(j)}|\boldsymbol{x})$ is 0, the second term $(II)$ is 0. Hence, $e^{-2\lambda T}(II)$ is 0. Similarly we can show that

$$\begin{aligned}
(III) =& 2\int_0^T dt \int_0^\infty d\tau'' \sum_{\boldsymbol{x}''\in\Omega} D_t\left(\tau'', \boldsymbol{x}''; a_0, \boldsymbol{x}_1\right) \\
&\left[\sum_{\boldsymbol{x}'''\in\Omega} \mathbb{T}_{\mathrm{F}}(\boldsymbol{x}'''|\boldsymbol{x}'') \int d\tau'\, \mathring{N}_{T-t}\left(\tau', \boldsymbol{x}; 0, \boldsymbol{x}'''\right)\right. \\
&\left.\sum_{\boldsymbol{x}_{l(i)}, \boldsymbol{x}_{r(i)}\in\Omega} g_{\boldsymbol{x}'\leftarrow\boldsymbol{x}}(\boldsymbol{x}_{l(i)}, \boldsymbol{x}_{r(i)})\mathbb{T}_{\mathrm{F}}(\boldsymbol{x}_{l(i)}|\boldsymbol{x})\mathbb{T}_{\mathrm{F}}(\boldsymbol{x}_{r(i)}|\boldsymbol{x})\right]^2.
\end{aligned} \tag{S6.59}$$

Since the sum $\sum_{\boldsymbol{x}_{l(i)}, \boldsymbol{x}_{r(i)}\in\Omega} g_{\boldsymbol{x}'\leftarrow\boldsymbol{x}}(\boldsymbol{x}_{l(i)}, \boldsymbol{x}_{r(i)})\mathbb{T}_{\mathrm{F}}(\boldsymbol{x}_{l(i)}|\boldsymbol{x})\mathbb{T}_{\mathrm{F}}(\boldsymbol{x}_{r(i)}|\boldsymbol{x})$ is 0, we also have $(III) = 0$. Hence, $e^{-2\lambda T}(III)$ is 0. This completes the proof of Eq. (S6.38). □

### S7 Identification of $\lambda$ from $\mathbb{T}_{\mathrm{F}}$ and $\pi_{\mathrm{B}}$

Here, we propose two methods to estimate the population growth rate $\lambda$. The first method is the direct calculation of $\lambda$ from the lineage tree by

$$\lambda = \frac{1}{T}\log\frac{N_T}{N_0}, \tag{S7.1}$$

where $T$ is the time length of the experiment, $N_T$ is the number of the cells at the end of the lineage tree, and $N_0$ is the number of the cells at the beginning of the experiment. In the analysis of E. coli data shown in Section 5 (C) of the main text, we calculated the $\lambda$ by a modification of this method because cells were flown from the chamber and $N_T$ is unknown in this case. We estimate $\lambda$ by

$$\lambda = \frac{1}{T}\frac{N}{C}, \tag{S7.2}$$

where $T$ is the time length of the experiment, $N$ is the number of the cells flown from the chamber, and $C$ is the average number of the cell in the chamber.

The second method is the analytical calculation from the estimated $\mathbb{T}_{\mathrm{F}}$ and $\pi_{\mathrm{B}}$. According to [14], the population growth rate $\lambda$ is given by the unique real number such that the largest eigen value of the matrix $M$ is unit. Recall that the matrix $M$ in the main (Eq. [5]) is defined by

$$M(\boldsymbol{x}'|\boldsymbol{x}) = \mathbb{T}_{\mathrm{F}}(\boldsymbol{x}'|\boldsymbol{x})Z(\boldsymbol{x}), \tag{S7.3}$$

where

$$Z(\boldsymbol{x}) = \int_0^\infty d\tau\, 2e^{-\lambda\tau}\pi_{\mathrm{F}}(\tau|\boldsymbol{x}). \tag{S7.4}$$

However, we cannot directly use this condition because the definition of $Z(\boldsymbol{x})$ is described by $\pi_{\mathrm{F}}$ while we only have $\mathbb{T}_{\mathrm{F}}$ and $\pi_{\mathrm{B}}$. Therefore, we need to calculate $Z(\boldsymbol{x})$ by using $\pi_{\mathrm{B}}$. By the definition of $\pi_{\mathrm{B}}$ (Eq. [3] in the main text), we obtain

$$\pi_{\mathrm{F}}(\tau|x) = \frac{Z(\boldsymbol{x})e^{\lambda\tau}\pi_{\mathrm{B}}(\tau|\boldsymbol{x})}{2}. \tag{S7.5}$$

Since $\pi_{\mathrm{F}}$ is normalized, we have

$$Z(\boldsymbol{x}) = \left(\int d\tau\, \frac{e^{\lambda\tau}\pi_{\mathrm{B}}(\tau|\boldsymbol{x})}{2}\right)^{-1}. \tag{S7.6}$$

By substituting $Z(\boldsymbol{x})$ in the definition of $M$, we have

$$M(\boldsymbol{x}'|\boldsymbol{x}) = \mathbb{T}_{\mathrm{F}}(\boldsymbol{x}'|\boldsymbol{x})\left(\int d\tau\, \frac{e^{\lambda\tau}\pi_{\mathrm{B}}(\tau|\boldsymbol{x})}{2}\right)^{-1}. \tag{S7.7}$$

Notice that this expression only contains $\mathbb{T}_{\mathrm{F}}$ and $\pi_{\mathrm{B}}$, which can be estimated by the EM algorithm.

Finally, we prove that $\lambda$ is uniquely determined by the above condition. The uniqueness is non-trivial because the forms of Eqs. (S7.3) and (S7.4) are different from those of Eqs. (S7.6) and (S7.7). Namely, we will show that there uniquely exists a real number $\alpha$ such that the largest eigenvalue of

$$M_\alpha(\boldsymbol{x}'|\boldsymbol{x}) = \mathbb{T}_{\mathrm{F}}(\boldsymbol{x}'|\boldsymbol{x})\left(\int d\tau\, \frac{e^{\alpha\tau}\pi_{\mathrm{B}}(\tau|\boldsymbol{x})}{2}\right)^{-1}, \tag{S7.8}$$

is unit. This proof is essentially the same as [14]. Let $\xi_\alpha$ be the largest eigenvalue of $M_\alpha$. The differentiation of $\xi_\alpha$ satisfies [20]

$$\frac{d\xi_\alpha}{d\alpha} = \frac{\sum_{\boldsymbol{x},\boldsymbol{x}'\in\Omega} u_\alpha(\boldsymbol{x}')\left(\frac{d}{d\alpha}M_\alpha(\boldsymbol{x}'|\boldsymbol{x})\right)v_\alpha(\boldsymbol{x})}{\sum_{\boldsymbol{x}\in\Omega} u_\alpha(\boldsymbol{x})v_\alpha(\boldsymbol{x})}, \tag{S7.9}$$

where $\boldsymbol{u}$ and $\boldsymbol{v}$ are the left and right eigenvectors of $M_\alpha$ corresponding to the largest eigenvalue, respectively. Since we assumed that $\mathbb{T}_{\mathrm{F}}$ is primitive, the matrix $M_\alpha$ is also primitive. By the Perron-Frobenius Theorem, $u_\alpha(\boldsymbol{x}) > 0$ and $v_\alpha(\boldsymbol{x}) > 0$ for all $\boldsymbol{x} \in \Omega$. By Eq. (S7.8), $\frac{d}{d\alpha}M_\alpha(\boldsymbol{x}'|\boldsymbol{x}) < 0$ for all $\boldsymbol{x}, \boldsymbol{x}' \in \Omega$. Therefore, $\frac{d\xi_\alpha}{d\alpha} < 0$. Also, we can easily see that $\lim_{\alpha\to\infty}\xi_\alpha = 0$ and $\lim_{\alpha\to-\infty}\xi_\alpha = \infty$. These results imply that $\xi_\alpha$ is monotonically decreasing from $\infty$ to zero with respect to $\alpha$. Hence, there uniquely exists $\alpha$ such that the largest eigenvalue of $M_\alpha$ is unit.

#### S8 Recursive computation of $\gamma_i$ and $\xi_{i,j}$ by belief propagation

We explain how to compute $\gamma_i$ and $\xi_{i,j}$ recursively by the Belief propagation. In the following, we use $\mathbb{P}(\boldsymbol{x})$ instead of $\mathbb{P}(\boldsymbol{x}_i = \boldsymbol{x})$ when the cell we focus on is clear from the context. Recall that $\gamma_i$ and $\xi_{i,j}$ are the posterior distributions defined as

$$\gamma_i(\boldsymbol{x}_i) = \mathbb{P}(\boldsymbol{x}_i|\mathcal{D}, \Theta^{(n)}), \tag{S8.1}$$
$$\xi_{i,j}(\boldsymbol{x}_i, \boldsymbol{x}_j) = \mathbb{P}(\boldsymbol{x}_i, \boldsymbol{x}_j|\mathcal{D}, \Theta^{(n)}), \tag{S8.2}$$

where $i$ and $j$ are the indices of a mother($i$) and daughter($j$) pair. $\Theta^{(n)}$ and $\mathcal{D}$ are the current parameter value and the observation data. First, we define two variables, $\alpha_i$ and $\beta_i$, as

$$\alpha_i(\boldsymbol{x}_i) = \mathbb{P}(\boldsymbol{x}_i, \mathrm{up}(i), \tau_i|\Theta^{(n)}), \tag{S8.3}$$
$$\beta_i(\boldsymbol{x}_i) = \mathbb{P}(\mathrm{down}(i)|\boldsymbol{x}_i, \Theta^{(n)}). \tag{S8.4}$$

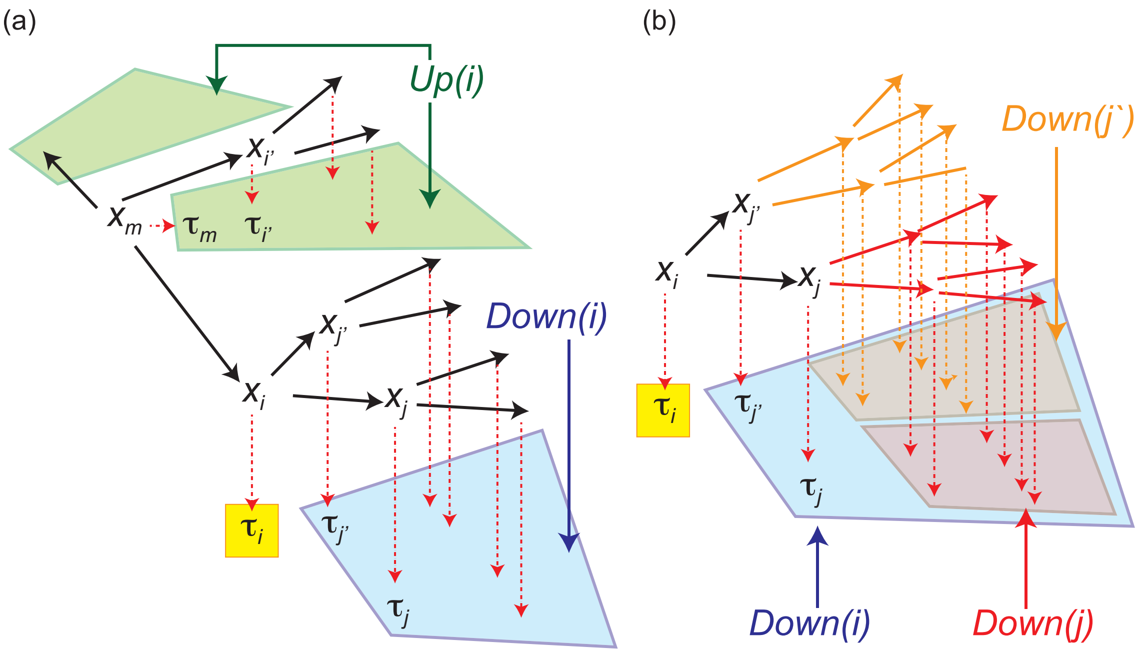


Figure S8.1: Graphical model representations of the relation among variables in a tree. (a) the relation among up($i$), down($i$), $\boldsymbol{x}_i$, and $\tau_i$. (b) The relations between $\boldsymbol{x}_j$, $\tau_j$, and down($j$) and $\boldsymbol{x}_{j'}$, $\tau_{j'}$, and down($j'$).

By focusing only on the cell $i$ in a lineage tree, $\mathcal{D}$ is decomposed into three parts: $\tau_i$, up($i$), and down($i$) as $\mathcal{D} = \mathrm{up}(i) \cup \{\tau_i\} \cup \mathrm{down}(i)$: down($i$) is the division time of the descendants of the cell $i$, and up($i$) is those of the other cells as $\mathrm{up}(i) := \mathcal{D} \setminus (\{\tau_i\} \cup \mathrm{down}(i))$ (Fig. S8.1(a)). By using this decomposition, $\gamma_i(\boldsymbol{x}_i)$ can be represented by $\alpha_i(\boldsymbol{x}_i)$ and $\beta_i(\boldsymbol{x}_i)$ as

$$\begin{aligned}\gamma_i(\boldsymbol{x}_i) &= \mathbb{P}(\boldsymbol{x}_i|\mathcal{D}, \Theta^{(n)}) = \mathbb{P}(\boldsymbol{x}_i, \mathcal{D}|\Theta^{(n)})/\mathbb{P}(\mathcal{D}|\Theta^{(n)}) \propto p(\boldsymbol{x}_i, \mathcal{D}|\Theta^{(n)}) \\ &= \mathbb{P}(\boldsymbol{x}_i, \mathrm{up}(i), \tau_i|\Theta^{(n)})\mathbb{P}(\mathrm{down}(i)|\boldsymbol{x}_i, \mathrm{up}(i), \tau_i, \Theta^{(n)}) \\ &= \mathbb{P}(\boldsymbol{x}_i, \mathrm{up}(i), \tau_i|\Theta^{(n)})\mathbb{P}(\mathrm{down}(i)|\boldsymbol{x}_i, \Theta^{(n)}) = \alpha_i(\boldsymbol{x}_i)\beta_i(\boldsymbol{x}_i),\end{aligned} \tag{S8.5}$$

where we used the fact that down($i$), up($i$), and $\{\tau_i\}$ are conditionally independent given $\boldsymbol{x}_i$ (Fig. S8.1(a)).

For $\xi_{ij}(\boldsymbol{x}_i, \boldsymbol{x}_j)$, we have

$$\xi_{i,j}(\boldsymbol{x}_i, \boldsymbol{x}_j) = \mathbb{P}(\boldsymbol{x}_i, \boldsymbol{x}_j|\mathcal{D}, \Theta^{(n)}) = \mathbb{P}(\boldsymbol{x}_i|\mathcal{D}, \Theta^{(n)})\mathbb{P}(\boldsymbol{x}_j|\boldsymbol{x}_i, \mathcal{D}, \Theta^{(n)}) = \gamma_i(\boldsymbol{x}_i)\mathbb{P}(\boldsymbol{x}_j|\boldsymbol{x}_i, \mathcal{D}, \Theta^{(n)}). \tag{S8.6}$$

For the factor $\mathbb{P}(\boldsymbol{x}_j|\boldsymbol{x}_i, \mathcal{D}, \Theta^{(n)})$, we have

$$\mathbb{P}(\boldsymbol{x}_j|\boldsymbol{x}_i, \mathcal{D}, \Theta^{(n)}) = \frac{\mathbb{P}(\boldsymbol{x}_j, \mathcal{D}|\boldsymbol{x}_i, \Theta^{(n)})}{\mathbb{P}(\mathcal{D}|\boldsymbol{x}_i, \Theta^{(n)})} = \frac{\mathbb{P}(\boldsymbol{x}_j, \mathcal{D}|\boldsymbol{x}_i, \Theta^{(n)})}{\sum_{\boldsymbol{x}_j \in \Omega} \mathbb{P}(\boldsymbol{x}_j, \mathcal{D}|\boldsymbol{x}_i, \Theta^{(n)})}. \tag{S8.7}$$

By focusing on the cells $j$, $\mathcal{D}$ can be decomposed as

$$\mathcal{D} = \mathrm{up}(j) \cup \{\tau_j\} \cup \mathrm{down}(j). \tag{S8.8}$$

By using this decomposition, we obtain

$$\begin{aligned}\mathbb{P}(\boldsymbol{x}_j, \mathcal{D}|\boldsymbol{x}_i, \Theta^{(n)}) &= \mathbb{P}(\mathrm{up}(j)|\boldsymbol{x}_i, \Theta^{(n)})\mathbb{P}(\boldsymbol{x}_j|\boldsymbol{x}_i, \mathrm{up}(j), \Theta^{(n)}) \\ &\quad \mathbb{P}(\tau_j|\boldsymbol{x}_i, \boldsymbol{x}_j, \mathrm{up}(j), \Theta^{(n)})\mathbb{P}(\mathrm{down}(j)|\boldsymbol{x}_i, \boldsymbol{x}_j, \tau_j, \mathrm{up}(j), \Theta^{(n)}) \\ &= \mathbb{P}(\mathrm{up}(j)|\boldsymbol{x}_i, \Theta^{(n)})\mathbb{P}(\boldsymbol{x}_j|\boldsymbol{x}_i, \Theta^{(n)})\mathbb{P}(\tau_j|\boldsymbol{x}_j, \Theta^{(n)})\mathbb{P}(\mathrm{down}(j)|\boldsymbol{x}_j, \Theta^{(n)}) \\ &= \mathbb{P}(\mathrm{up}(j)|\boldsymbol{x}_i, \Theta^{(n)})\mathbb{T}_{\mathrm{F}}(\boldsymbol{x}_j|\boldsymbol{x}_i)\pi_{\mathrm{B}}(\tau_j \mid \boldsymbol{x}_j)\beta_j(\boldsymbol{x}_j),\end{aligned} \tag{S8.9}$$

here and hereafter we abbreviate $\mathbb{T}_{\mathrm{F}}(\boldsymbol{x}_j|\boldsymbol{x}_i, \Theta^{(n)})$ and $\pi_{\mathrm{B}}(\tau_j|\boldsymbol{x}_j, \Theta^{(n)})$ with $\mathbb{T}_{\mathrm{F}}(\boldsymbol{x}_j|\boldsymbol{x}_i)$ and $\pi_{\mathrm{B}}(\tau_j|\boldsymbol{x}_j)$. Since $\mathbb{P}(\mathrm{up}(j)|\boldsymbol{x}_i, \Theta^{(n)})$ is constant with respect to $\boldsymbol{x}_j$, we have

$$\xi_{i,j}(\boldsymbol{x}_i, \boldsymbol{x}_j) = \gamma_i(\boldsymbol{x}_i)\mathbb{P}(\boldsymbol{x}_j|\boldsymbol{x}_i, \mathcal{D}, \Theta^{(n)}) = \gamma_i(\boldsymbol{x}_i)\frac{\mathbb{T}_{\mathrm{F}}(\boldsymbol{x}_j|\boldsymbol{x}_i)\pi_{\mathrm{B}}(\tau_j \mid \boldsymbol{x}_j)\beta_j(\boldsymbol{x}_j)}{\sum_{\boldsymbol{x}_j} \mathbb{T}_{\mathrm{F}}(\boldsymbol{x}_j|\boldsymbol{x}_i)\pi_{\mathrm{B}}(\tau_j \mid \boldsymbol{x}_j)\beta_j(\boldsymbol{x}_j)}. \tag{S8.10}$$

Because both $\gamma_i(\boldsymbol{x}_i)$ and $\xi_{i,j}(\boldsymbol{x}_i, \boldsymbol{x}_j)$ are described as functions of $\alpha_i(\boldsymbol{x}_i)$ and $\beta_i(\boldsymbol{x}_i)$, the remaining task is to compute $\alpha_i(\boldsymbol{x}_i)$ and $\beta_i(\boldsymbol{x}_i)$, recursively. We first derive a recursive formula of $\beta_i(\cdot)$. Let $j$ and $j'$ be the indices of the daughters of the cell $i$. Then, $\mathrm{down}(i)$ is decomposed as $\mathrm{down}(i) = \{\tau_j\} \cup \{\tau_{j'}\} \cup \mathrm{down}(j) \cup \mathrm{down}(j')$ (Fig. S8.1(b)). Since $\{\tau_j\}$, $\{\tau_{j'}\}$, $\mathrm{down}(j)$, and $\mathrm{down}(j')$ are conditionally independent given $\boldsymbol{x}_j$ and $\boldsymbol{x}_{j'}$, we have

$$\begin{aligned}\beta_i(\boldsymbol{x}_i) &= \sum_{\boldsymbol{x}_j, \boldsymbol{x}_{j'}} \mathbb{P}(\boldsymbol{x}_j, \boldsymbol{x}_{j'}, \mathrm{down}(i)|\boldsymbol{x}_i, \Theta^{(n)}) \\ &= \sum_{\boldsymbol{x}_j, \boldsymbol{x}_{j'}} \mathbb{P}(\boldsymbol{x}_j|\boldsymbol{x}_i, \Theta^{(n)})\mathbb{P}(\boldsymbol{x}_{j'}|\boldsymbol{x}_i, \Theta^{(n)})\mathbb{P}(\mathrm{down}(j), \tau_j|\boldsymbol{x}_j, \Theta^{(n)})\mathbb{P}(\mathrm{down}(j'), \tau_{j'}|\boldsymbol{x}_{j'}, \Theta^{(n)}) \\ &= \sum_{\boldsymbol{x}_j, \boldsymbol{x}_{j'}} \mathbb{T}_{\mathrm{F}}(\boldsymbol{x}_j|\boldsymbol{x}_i)\mathbb{T}_{\mathrm{F}}(\boldsymbol{x}_{j'}|\boldsymbol{x}_i)\mathbb{P}(\mathrm{down}(j)|\boldsymbol{x}_j, \Theta^{(n)})\mathbb{P}(\mathrm{down}(j')|\boldsymbol{x}_{j'}, \Theta^{(n)})\pi_{\mathrm{B}}(\tau_j|\boldsymbol{x}_j)\pi_{\mathrm{B}}(\tau_{j'}|\boldsymbol{x}_{j'}) \\ &= \left[\sum_{\boldsymbol{x}_j} \pi_{\mathrm{B}}(\tau_j|\boldsymbol{x}_j)\mathbb{T}_{\mathrm{F}}(\boldsymbol{x}_j|\boldsymbol{x}_i)\beta_j(\boldsymbol{x}_j)\right]\left[\sum_{\boldsymbol{x}_{j'}} \pi_{\mathrm{B}}(\tau_{j'}|\boldsymbol{x}_{j'})\mathbb{T}_{\mathrm{F}}(\boldsymbol{x}_{j'}|\boldsymbol{x}_i)\beta_{j'}(\boldsymbol{x}_{j'})\right].\end{aligned} \tag{S8.11}$$

At the initial step of the recursion, $\beta_i(\cdot)$, for the non-leaf cells at the end of the lineage tree, are set as $\beta_i(\cdot) = 1$. Then the other $\beta_i(\cdot)$s are calculated by back propagating the values of $\beta_i(\cdot)$s from the leaves toward the root. We next derive a recursive formula of $\alpha_i(\cdot)$. Let $m$ be the index of the mother of cell $i$, and $i'$ be the index of the sibling of the cell $i$. Using the conditional independence of $\mathrm{up}(i)$ and $\{x_i, \tau_i\}$ given $\boldsymbol{x}_m$ (Fig. S8.1(a)), we obtain

$$\alpha_i(\boldsymbol{x}_i) = \sum_{\boldsymbol{x}_m} \mathbb{P}(\boldsymbol{x}_m, \boldsymbol{x}_i, \mathrm{up}(i), \tau_i|\Theta^{(n)}) = \sum_{\boldsymbol{x}_m} \mathbb{P}(\boldsymbol{x}_m, \mathrm{up}(i)|\Theta^{(n)})\mathbb{T}_{\mathrm{F}}(\boldsymbol{x}_i|\boldsymbol{x}_m)\pi_{\mathrm{B}}(\tau_i|\boldsymbol{x}_i). \tag{S8.12}$$

By using a decomposition of $\mathrm{up}(i)$ as

$$\mathrm{up}(i) = \{\tau_m\} \cup \mathrm{up}(m) \cup \{\tau_{i'}\} \cup \mathrm{down}(i'), \tag{S8.13}$$

$\mathbb{P}(\boldsymbol{x}_m, \mathrm{up}(i)|\Theta^{(n)})$ becomes

$$\begin{aligned}\mathbb{P}(\boldsymbol{x}_m, \mathrm{up}(i)|\Theta^{(n)}) &= \sum_{\boldsymbol{x}_{i'}} \mathbb{P}(\boldsymbol{x}_m, \boldsymbol{x}_{i'}, \tau_m, \mathrm{up}(m), \tau_{i'}, \mathrm{down}(i')|\Theta^{(n)}) \\ &= \sum_{\boldsymbol{x}_{i'}} \mathbb{P}(\boldsymbol{x}_m, \mathrm{up}(m), \tau_m|\Theta^{(n)})\mathbb{T}_{\mathrm{F}}(\boldsymbol{x}_{i'}|\boldsymbol{x}_m)\pi_{\mathrm{B}}(\tau_{i'}|\boldsymbol{x}_{i'})\mathbb{P}(\mathrm{down}(i')|\boldsymbol{x}_{i'}, \Theta^{(n)}) \\ &= \sum_{\boldsymbol{x}_{i'}} \alpha_m(\boldsymbol{x}_m)\mathbb{T}_{\mathrm{F}}(\boldsymbol{x}_{i'}|\boldsymbol{x}_m)\pi_{\mathrm{B}}(\tau_{i'}|\boldsymbol{x}_{i'})\beta_{i'}(\boldsymbol{x}_{i'}).\end{aligned} \tag{S8.14}$$

Combining these two equations, we have

$$\alpha_i(\boldsymbol{x}_i) = \sum_{\boldsymbol{x}_m} \left[ \sum_{\boldsymbol{x}_{i'}} \alpha_m(\boldsymbol{x}_m) \mathbb{T}_{\mathrm{F}}(\boldsymbol{x}_{i'}|\boldsymbol{x}_m) \pi_{\mathrm{B}}(\tau_{i'}|\boldsymbol{x}_{i'}) \beta_{i'}(\boldsymbol{x}_{i'}) \right] \mathbb{T}_{\mathrm{F}}(\boldsymbol{x}_i|\boldsymbol{x}_m) \pi_{\mathrm{B}}(\tau_i|\boldsymbol{x}_i). \tag{S8.15}$$

If the number of the daughters is not necessarily two, then the inner summation of the above expression (enclosed by brackets) appears more for all the labels of the daughters $i', i'', \ldots$ other than the cell $i$. For example, if the cell $m$ has two daughters labeled by $i', i''$ other than the cell $i$, then Eq. (S8.15) becomes

$$\alpha_i(\boldsymbol{x}_i) = \sum_{\boldsymbol{x}_m} \left[ \sum_{\boldsymbol{x}_{i'}} \alpha_m(\boldsymbol{x}_m) \mathbb{T}_{\mathrm{F}}(\boldsymbol{x}_{i'}|\boldsymbol{x}_m) \pi_{\mathrm{B}}(\tau_{i'}|\boldsymbol{x}_{i'}) \beta_{i'}(\boldsymbol{x}_{i'}) \right] \left[ \sum_{\boldsymbol{x}_{i''}} \alpha_m(\boldsymbol{x}_m) \mathbb{T}_{\mathrm{F}}(\boldsymbol{x}_{i''}|\boldsymbol{x}_m) \pi_{\mathrm{B}}(\tau_{i''}|\boldsymbol{x}_{i''}) \beta_{i''}(\boldsymbol{x}_{i''}) \right] \mathbb{T}_{\mathrm{F}}(\boldsymbol{x}_i|\boldsymbol{x}_m) \pi_{\mathrm{B}}(\tau_i|\boldsymbol{x}_i). \tag{S8.16}$$

If the cell $i$ is the only daughter of the cell $m$, then the inner summation is substituted by one. At the initial step of the recursion, $\alpha_{\mathrm{root}}(\boldsymbol{x}_{\mathrm{root}})$ for the root cell is calculated as

$$\alpha_{\mathrm{root}}(\boldsymbol{x}_{\mathrm{root}}) = \rho(\boldsymbol{x}_{\mathrm{root}}) \pi_{\mathrm{B}}(\tau_{\mathrm{root}}|\boldsymbol{x}_{\mathrm{root}}), \tag{S8.17}$$

where $\rho(\boldsymbol{x}_{\mathrm{root}})$ is the distribution of the state of the root cell. If we can observe the stationary distribution of the population of the cells, we can use that distribution for $\rho(\boldsymbol{x}_{\mathrm{root}})$ by presuming that the root cell is a sample from the stationary distribution. If we do not have $\rho(\boldsymbol{x}_{\mathrm{root}})$, we can estimate $\rho(x)$ as $\beta_{\mathrm{root}}(x)$ by MLE.

### S9 Update of the parameters in M-step

In this section, we explain the detail on the update of the parameters in the M-step of the BW algorithm. Before explaining the detail, we first review the sufficient statistics of the exponential family. Each probability distribution in the exponential family has its sufficient statistics, which is the key tool to find the maximum likelihood estimator. Suppose that we have data $\{\boldsymbol{x}_1, \boldsymbol{x}_2, \ldots, \boldsymbol{x}_N\}$ and want to compute the maximum likelihood estimator of a distribution $f(\cdot|\boldsymbol{\theta})$ that belongs to the exponential family with a vector of parameters $\boldsymbol{\theta}$. Let $\boldsymbol{\eta}(\boldsymbol{x})$ be the sufficient statistics of the distribution and $\boldsymbol{\eta}[\boldsymbol{\theta}] = \int d\boldsymbol{x}\, \boldsymbol{\eta}(\boldsymbol{x}) f(\boldsymbol{x}|\boldsymbol{\theta})$. Then, the maximum likelihood estimator is computed as the solution of the moment-matching equation [21]:

$$\boldsymbol{\eta}[\boldsymbol{\theta}] = \int d\boldsymbol{x}\, \boldsymbol{\eta}(\boldsymbol{x}) f_{\mathrm{emp}}(\boldsymbol{x}) = \frac{1}{N} \sum_{i=1}^{N} \boldsymbol{\eta}(\boldsymbol{x}_i), \tag{S9.1}$$

where the empirical distribution is defined by

$$f_{\mathrm{emp}}(\boldsymbol{x}) := \frac{1}{N} \sum_{i=1}^{N} \delta(\boldsymbol{x} - \boldsymbol{x}_i). \tag{S9.2}$$

The left-hand-side of (S9.1) is determined by the model and the parameter $\boldsymbol{\theta}$, and the right-hand-side is determined by data. We choose the parameter vector $\boldsymbol{\theta}$ so that the left-hand-side coincides with the right-hand-side.

In the M-step, the parameters $\Theta^{(n)} = (\{\boldsymbol{\theta}^B_{\boldsymbol{x}}\}_{\boldsymbol{x}\in\Omega}, \mathbb{T}_{\mathrm{F}})$ are updated so that $\pi_{\mathrm{B}}(\cdot|\boldsymbol{\theta}^B_{\boldsymbol{x}})$ and $\mathbb{T}_{\mathrm{F}}$ fit Eq. (S5.11), i.e., we choose the parameters that maximize the likelihood weighted by the posterior distributions $\gamma_i(\boldsymbol{x})$ and $\xi_{i,j}(\boldsymbol{x}, \boldsymbol{x}')$. We first compute $\boldsymbol{\theta}^B_{\boldsymbol{x}}$. After the E-step, we obtain an approximation of the empirical distribution of the division time of the cells with state $\boldsymbol{x}$ as

$$\pi^{\mathrm{BW}}_{\mathrm{emp}}(\tau|\boldsymbol{x}) = \frac{1}{\sum_i \gamma_i(\boldsymbol{x})} \sum_{i\in\mathcal{T}} \gamma_i(\boldsymbol{x}) \delta(\tau - \tau_i). \tag{S9.3}$$

By using this approximated empirical distribution instead of the true empirical distribution in Eq. (S9.1), we have

$$\boldsymbol{\eta}_B[\boldsymbol{\theta}_{\boldsymbol{x}}^B] = \int d\tau\, \boldsymbol{\eta}_B(\tau)\pi_{\mathrm{emp}}^{\mathrm{BW}}(\tau|\boldsymbol{x}) = \frac{1}{\sum_i \gamma_i(\boldsymbol{x})}\sum_{i\in\mathcal{T}}\gamma_i(\boldsymbol{x})\boldsymbol{\eta}_B(\tau_i), \tag{S9.4}$$

where $\boldsymbol{\eta}_B$ is the sufficient statistics of $\pi_{\mathrm{B}}$ We next compute the transition matrix. The transition matrix is updated so that $\mathbb{T}_{\mathrm{F}}(\boldsymbol{x}|\boldsymbol{x}')$ becomes proportional to $\sum_{i,j}\xi_{i,j}(\boldsymbol{x},\boldsymbol{x}')$, where the pair indices $(i,j)$ runs over all parent-daughter pairs.

### S10 Recursive computation of $\gamma_i$ and $\xi_{i,j}$ for the continuous-state model

The recursive computation explained in Section S8 is still applicable when $\Omega$ is continuous. For the continuous model with the linear dynamics,

$$\boldsymbol{x}' = \boldsymbol{A}\boldsymbol{x} + \boldsymbol{w}, \tag{S10.1}$$

$$\log\tau = \boldsymbol{C}\boldsymbol{x} + v, \tag{S10.2}$$

we can use a different recursive computation developed for the Kalman smoother, which exploits the unique property of the normal distributions that the conversion of a normal distribution either by linear noisy dynamics or by the Bayes' rule generates another normal distribution and thereby that the conversions can be represented by updates of the sufficient statistics of the normal distributions, i.e., their means and covariances. In this setting, $\Theta^{(n)}$ includes the entities of $\boldsymbol{A}$, $\boldsymbol{C}$, $\boldsymbol{\Sigma}_{\boldsymbol{w}}$ and $\Sigma_v$ after the $k$th iteration. Here, we explain how the recursion of the Kalman smoother can be extended to the tree-structured model, which was developed in [22].

In order to derive the recursions of $\gamma_i(\boldsymbol{x}_i)$ and $\xi_{i,j}(\boldsymbol{x}_i,\boldsymbol{x}_j)$ as updates of the means and the variances of normal distributions, $\beta_i(\boldsymbol{x}_i) := \mathbb{P}(\mathrm{down}(i)|\boldsymbol{x}_i,\Theta^{(n)})$ used in the previous section is not appropriate, because $\beta_i(\boldsymbol{x}_i)$ is not a distribution of $\boldsymbol{x}_i$. Therefore, we employ another recursive computation formulae of $\gamma_i(\boldsymbol{x}_i)$ and $\xi_{i,j}(\boldsymbol{x}_i,\boldsymbol{x}_j)$, which are fit for this purpose.

Suppose that the cell $j$ is a daughter of the cell $i$, then $\gamma_j(\boldsymbol{x}_j)$ for the daughter is decomposed as

$$\gamma_j(\boldsymbol{x}_j) := \mathbb{P}(\boldsymbol{x}_j|\mathcal{D},\Theta^{(n)}) = \sum_{\boldsymbol{x}_i}\mathbb{P}(\boldsymbol{x}_i,\boldsymbol{x}_j|\mathcal{D}) = \sum_{\boldsymbol{x}_i}\mathbb{P}(\boldsymbol{x}_j|\boldsymbol{x}_i,\mathcal{D})\mathbb{P}(\boldsymbol{x}_i|\mathcal{D}) = \sum_{\boldsymbol{x}_i}\mathbb{P}(\boldsymbol{x}_j|\boldsymbol{x}_i,\mathcal{D})\gamma_i(\boldsymbol{x}_i), \tag{S10.3}$$

where and hereafter we abbreviate the dependencies of probabilities on the parameter $\Theta^{(n)}$ for the notational simplicity. Similarly, $\xi_{i,j}(\boldsymbol{x}_i,\boldsymbol{x}_j)$ is decomposed as

$$\xi_{i,j}(\boldsymbol{x}_i,\boldsymbol{x}_j) = \gamma_i(\boldsymbol{x}_i)\mathbb{P}(\boldsymbol{x}_j|\boldsymbol{x}_i,\mathcal{D}). \tag{S10.4}$$

$\mathbb{P}(\boldsymbol{x}_j|\boldsymbol{x}_i,\mathcal{D})$ is calculated as

$$\mathbb{P}(\boldsymbol{x}_j|\boldsymbol{x}_i,\mathcal{D}) = \mathbb{P}(\boldsymbol{x}_j|\boldsymbol{x}_i,\tau_j,\mathrm{down}(j)) = \frac{\mathbb{P}(\boldsymbol{x}_i|\boldsymbol{x}_j)\mathbb{P}(\boldsymbol{x}_j|\tau_j,\mathrm{down}(j)))}{\mathbb{P}(\boldsymbol{x}_i|\tau_j,\mathrm{down}(j))} \propto \mathbb{P}(\boldsymbol{x}_i|\boldsymbol{x}_j)\bar{\beta}_j(\boldsymbol{x}_j). \tag{S10.5}$$

where we define $\bar{\beta}_i(\boldsymbol{x}_i)$ as

$$\bar{\beta}_i(\boldsymbol{x}_i) := \mathbb{P}(\boldsymbol{x}_i|\mathrm{down}(i),\tau_i,\Theta^{(n)}). \tag{S10.6}$$

Note that $\bar{\beta}_i(\boldsymbol{x}_i)$ defined here is related to $\beta_i(\boldsymbol{x}_i) := \mathbb{P}(\mathrm{down}(i)|\boldsymbol{x}_i,\Theta^{(n)})$ as $\bar{\beta}_i(\boldsymbol{x}_i) = \pi_{\mathrm{B}}(\tau_i|\boldsymbol{x}_i)\beta_i(\boldsymbol{x}_i)\mathbb{P}(\boldsymbol{x}_i)$:

$$\bar{\beta}_i(\boldsymbol{x}_i) = \frac{\mathbb{P}(\boldsymbol{x}_i,\tau_i,\mathrm{down}(i))}{\mathbb{P}(\tau_i,\mathrm{down}(i))} = \frac{\mathbb{P}(\tau_i|\boldsymbol{x}_i,\mathrm{down}(i))\mathbb{P}(\mathrm{down}(i)|\boldsymbol{x}_i)\mathbb{P}(\boldsymbol{x}_i)}{\mathbb{P}(\tau_i,\mathrm{down}(i))} \propto \pi_{\mathrm{B}}(\tau_i|\boldsymbol{x}_i)\beta_i(\boldsymbol{x}_i)\mathbb{P}(\boldsymbol{x}_i). \tag{S10.7}$$

$\mathbb{P}(\boldsymbol{x}_i|\boldsymbol{x}_j)$ in Eq. (S10.5) is obtained by Bayes's theorem as

$$\mathbb{P}(\boldsymbol{x}_i|\boldsymbol{x}_j) = \frac{\mathbb{T}_{\mathrm{F}}(\boldsymbol{x}_j|\boldsymbol{x}_i)}{\mathbb{P}(\boldsymbol{x}_j)}\mathbb{P}(\boldsymbol{x}_i). \tag{S10.8}$$

Among the terms in Eq. (S10.5), $\mathbb{P}(\boldsymbol{x}_j)$ is the unconditioned prior distribution and can be computed as

$$\mathbb{P}(\boldsymbol{x}_j) = \sum_{\boldsymbol{x}_i} \mathbb{T}_{\mathrm{F}}(\boldsymbol{x}_j|\boldsymbol{x}_i)\mathbb{P}(\boldsymbol{x}_i). \tag{S10.9}$$

Starting from the prior distribution of the state of the root cell in the tree, $\mathbb{P}(\boldsymbol{x}_i)$ is obtained by repeating this calculation down to $\mathbb{P}(\boldsymbol{x}_i)$. Thus $\mathbb{P}(\boldsymbol{x}_j|\boldsymbol{x}_i, \mathcal{D})$ is also computable as long as $\bar{\beta}_i(\boldsymbol{x}_i)$ is available. $\bar{\beta}_i(\boldsymbol{x}_i)$ is converted by Bayes' rule as

$$\begin{aligned}\bar{\beta}_i(\boldsymbol{x}_i) &= \frac{\mathbb{P}(\boldsymbol{x}_i, \tau_i|\mathrm{down}(i), \Theta^{(n)})}{\mathbb{P}(\tau_i|\mathrm{down}(i))} = \frac{\mathbb{P}(\tau_i|\boldsymbol{x}_i, \mathrm{down}(i))\mathbb{P}(\boldsymbol{x}_i|\mathrm{down}(i))}{\mathbb{P}(\tau_i|\mathrm{down}(i))} \\ &= \frac{\mathbb{P}(\tau_i|\boldsymbol{x}_i)\mathbb{P}(\boldsymbol{x}_i|\mathrm{down}(i))}{\mathbb{P}(\tau_i|\mathrm{down}(i))} = \frac{\pi_{\mathrm{B}}(\tau_i|\boldsymbol{x}_i)\mathbb{P}(\boldsymbol{x}_i|\mathrm{down}(i))}{\mathbb{P}(\tau_i|\mathrm{down}(i))}.\end{aligned} \tag{S10.10}$$

Let $j'$ be the other daughter of the cell $i$ than the cell $j$. Since $\mathrm{down}(i)$ is then decomposed as $\mathrm{down}(j) \cup \{\tau_j\} \cup \mathrm{down}(j') \cup \{\tau_{j'}\}$, we obtain

$$\begin{aligned}\mathbb{P}(\boldsymbol{x}_i|\mathrm{down}(i)) &= \frac{\mathbb{P}(\mathrm{down}(i)|\boldsymbol{x}_i)\mathbb{P}(\boldsymbol{x}_i)}{\mathbb{P}(\mathrm{down}(i))} = \frac{\mathbb{P}(\mathrm{down}(j), \tau_j, \mathrm{down}(j'), \tau_{j'}|\boldsymbol{x}_i)\mathbb{P}(\boldsymbol{x}_i)}{\mathbb{P}(\mathrm{down}(i))} \\ &= \frac{\mathbb{P}(\mathrm{down}(j), \tau_j|\boldsymbol{x}_i)\mathbb{P}(\mathrm{down}(j'), \tau_{j'}|\boldsymbol{x}_i)}{\mathbb{P}(\mathrm{down}(i))}\mathbb{P}(\boldsymbol{x}_i),\end{aligned} \tag{S10.11}$$

where we use the conditional independence of $\{\mathrm{down}(j), \tau_j\}$ and $\{\mathrm{down}(j'), \tau_{j'}\}$ given $\boldsymbol{x}_j$ at the second line. By applying Bayes's theorem to $\mathbb{P}(\mathrm{down}(j), \tau_j|\boldsymbol{x}_i)$ and $\mathbb{P}(\mathrm{down}(j'), \tau_{j'}|\boldsymbol{x}_i)$, we have

$$\begin{aligned}\mathbb{P}(\boldsymbol{x}_i|\mathrm{down}(i)) &= \frac{\mathbb{P}(\mathrm{down}(j), \tau_j)\mathbb{P}(\boldsymbol{x}_i|\mathrm{down}(j), \tau_j)\mathbb{P}(\mathrm{down}(j'), \tau_{j'})\mathbb{P}(\boldsymbol{x}_i|\mathrm{down}(j'), \tau_{j'})}{\mathbb{P}(\mathrm{down}(i))}\mathbb{P}(\boldsymbol{x}_i)^{-1} \\ &\propto \mathbb{P}(x_i|\mathrm{down}(j), \tau_j)\mathbb{P}(\boldsymbol{x}_i|\mathrm{down}(j'), \tau_{j'})\mathbb{P}(\boldsymbol{x}_i)^{-1} \\ &= \bar{\beta}_{j\to i}(\boldsymbol{x}_i)\bar{\beta}_{j'\to i}(\boldsymbol{x}_i)\mathbb{P}(\boldsymbol{x}_i)^{-1},\end{aligned} \tag{S10.12}$$

where we define

$$\bar{\beta}_{j\to i}(\boldsymbol{x}_i) := \mathbb{P}(\boldsymbol{x}_i|\tau_j, \mathrm{down}(j), \Theta^{(n)}). \tag{S10.13}$$

Finally, $\bar{\beta}_{j\to i}(\boldsymbol{x}_i)$ is computed recursively as follows:

$$\begin{aligned}\bar{\beta}_{j\to i}(\boldsymbol{x}_i) &= \int d\boldsymbol{x}_j \mathbb{P}(\boldsymbol{x}_i, \boldsymbol{x}_j|\tau_j, \mathrm{down}(j)) = \int d\boldsymbol{x}_j \mathbb{P}(\boldsymbol{x}_i|\boldsymbol{x}_j, \tau_j, \mathrm{down}(j))\mathbb{P}(\boldsymbol{x}_j|\tau_j, \mathrm{down}(j)) \\ &= \int d\boldsymbol{x}_j \mathbb{P}(\boldsymbol{x}_i|\boldsymbol{x}_j)\bar{\beta}_j(\boldsymbol{x}_j),\end{aligned} \tag{S10.14}$$

where we use Eq. (S10.8).

In summary, $\bar{\beta}_i(\boldsymbol{x}_i)$ is obtained given $\bar{\beta}_j(\boldsymbol{x}_j)$, $\bar{\beta}_{j'}(\boldsymbol{x}_j)$, $\mathbb{P}(\boldsymbol{x}_i)$, $\mathbb{T}_{\mathrm{F}}(\boldsymbol{x}_j|\boldsymbol{x}_i)$, and $\pi_{\mathrm{B}}(\tau_i|\boldsymbol{x}_i)$ as follows:

From Eq. (S10.9),
$$\mathbb{P}(\boldsymbol{x}_j) = \sum_{\boldsymbol{x}_j} \mathbb{T}_{\mathrm{F}}(\boldsymbol{x}_j|\boldsymbol{x}_i)\mathbb{P}(\boldsymbol{x}_i) \tag{S10.15}$$

From Eq. (S10.8),
$$\mathbb{P}(\boldsymbol{x}_i|\boldsymbol{x}_j) = \frac{\mathbb{T}_{\mathrm{F}}(\boldsymbol{x}_j|\boldsymbol{x}_i)}{\mathbb{P}(\boldsymbol{x}_j)}\mathbb{P}(\boldsymbol{x}_i), \tag{S10.16}$$

From Eq. (S10.14),
$$\bar{\beta}_{j\to i}(\boldsymbol{x}_i) = \int d\boldsymbol{x}_j \mathbb{P}(\boldsymbol{x}_i|\boldsymbol{x}_j)\bar{\beta}_j(\boldsymbol{x}_j), \tag{S10.17}$$

From Eq. (S10.12)
$$\mathbb{P}(\boldsymbol{x}_i|\mathrm{down}(i)) = \bar{\beta}_{j\to i}(\boldsymbol{x}_i)\bar{\beta}_{j'\to i}(\boldsymbol{x}_i)\mathbb{P}(\boldsymbol{x}_i)^{-1}, \tag{S10.18}$$

From Eq. (S10.10),
$$\bar{\beta}_i(\boldsymbol{x}_i) \propto \pi_{\mathrm{B}}(\tau_i|\boldsymbol{x}_i)\mathbb{P}(\boldsymbol{x}_i|\mathrm{down}(i)). \tag{S10.19}$$

In the case where the cell $i$ has $z$ daughters labeled by $j_1, j_2, \ldots, j_z$, Eq. (S10.18) is replaced by

$$\mathbb{P}(\boldsymbol{x}_i|\mathrm{down}(i)) = \left[\prod_{m=1}^{z} \bar{\beta}_{j_m\to i}(\boldsymbol{x}_i)\right]\mathbb{P}(\boldsymbol{x}_i)^{1-z}. \tag{S10.20}$$

This equation is derived by a similar argument. At the end of the lineages in the tree, $\bar{\beta}_i$ is initialized as

$$\bar{\beta}_i(\boldsymbol{x}_i) = \mathbb{P}(\boldsymbol{x}_i|\tau_i) = \frac{\mathbb{P}(\boldsymbol{x}_i, \tau_i)}{\mathbb{P}(\tau_i)} \propto \pi_{\mathrm{B}}(\tau_i|\boldsymbol{x}_i)\mathbb{P}(\boldsymbol{x}_i). \tag{S10.21}$$

Then, $\gamma_j(\boldsymbol{x}_j)$ and $\xi_{i,j}(\boldsymbol{x}_i, \boldsymbol{x}_j)$ are obtained from $\bar{\beta}_i(\boldsymbol{x}_i)$ via $\mathbb{P}(\boldsymbol{x}_j|\boldsymbol{x}_i, \mathcal{D})$ as

From Eq. (S10.5), $$\mathbb{P}(\boldsymbol{x}_j|\boldsymbol{x}_i, \mathcal{D}) \propto \mathbb{P}(\boldsymbol{x}_i|\boldsymbol{x}_j)\bar{\beta}_j(\boldsymbol{x}_j), \tag{S10.22}$$

From Eq. (S10.3), $$\gamma_j(\boldsymbol{x}_j) = \sum_{\boldsymbol{x}_i} \mathbb{P}(\boldsymbol{x}_j|\boldsymbol{x}_i, \mathcal{D})\gamma_i(\boldsymbol{x}_i), \tag{S10.23}$$

From Eq. (S10.4), $$\xi_{i,j}(\boldsymbol{x}_i, \boldsymbol{x}_j) = \gamma_i(\boldsymbol{x}_i)\mathbb{P}(\boldsymbol{x}_j|\boldsymbol{x}_i, \mathcal{D}). \tag{S10.24}$$

It should be noted that the linearity of the underlying dynamics, i.e., Eq. (S10.1), is not assumed up to this point. Thereby, these recursive formulae hold for general continuous state models.

We next convert these formulae into the form of the update rules of the means and the covariances of the corresponding normal distributions by assuming Eq. (S10.1) for dynamics. The prior probability of the state of the root cell is assumed to follow a multivariate normal distribution with the mean vector $\boldsymbol{\mu}_{\mathbf{root}}$ and the covariance matrix $\boldsymbol{\Sigma}_{\mathbf{root}}$. Then, because Eq. (S10.1) is a linear dynamics with additive noises, $\mathbb{P}(\boldsymbol{x}_i)$ is also a normal distribution with the mean vector $\boldsymbol{\mu}_i^{\mathrm{p}}$ and the covariance matrix $\boldsymbol{\Sigma}_i^{\mathrm{p}}$. Then, Eq. (S10.15) is represented as

$$\boldsymbol{\mu}_j^{\mathrm{p}} = \boldsymbol{A}\boldsymbol{\mu}_i^{\mathrm{p}}, \qquad \boldsymbol{\Sigma}_j^{\mathrm{p}} = \boldsymbol{\Sigma}_{\boldsymbol{w}} + \boldsymbol{A}^{\mathbb{T}}\boldsymbol{\Sigma}_i^{\mathrm{p}}\boldsymbol{A}. \tag{S10.25}$$

The root cell is initialized as $\boldsymbol{\mu}_{\mathrm{root}}^{\mathrm{p}} = \mu_{\mathrm{root}}$ and $\boldsymbol{\Sigma}_{\mathrm{root}}^{\mathrm{p}} = \Sigma_{\mathrm{root}}$. Similarly, $\mathbb{P}(\boldsymbol{x}_i|\boldsymbol{x}_j)$ is a multivariate normal distribution with the parameter $\{\boldsymbol{\mu}_{i|j}^{\mathrm{Bay}}, \boldsymbol{\Sigma}_{i|j}^{\mathrm{Bay}}\}$ and Eq. (S10.16) is represented as

$$\boldsymbol{\Sigma}_{i|j}^{\mathrm{Bay}} = ((\boldsymbol{\Sigma}_i^{\mathrm{p}})^{-1} + \boldsymbol{A}^{\mathbb{T}}\boldsymbol{\Sigma}_{\boldsymbol{w}}^{-\mathbf{1}}\boldsymbol{A})^{-1}, \qquad \boldsymbol{\mu}_{i|j}^{\mathrm{Bay}} = \boldsymbol{\Sigma}_{i|j}^{\mathrm{Bay}}(\boldsymbol{A}^{\mathbb{T}}\boldsymbol{\Sigma}_{\boldsymbol{w}}^{-1}\boldsymbol{x}_j + (\boldsymbol{\Sigma}_i^{\mathrm{p}})^{-1}\boldsymbol{\mu}_i^{\mathrm{p}}). \tag{S10.26}$$

Let the parameters of $\bar{\beta}_i(\boldsymbol{x}_i)$ and $\bar{\beta}_{j\to i}(\boldsymbol{x}_i)$ be $\{\boldsymbol{\mu}_i^{\beta}, \boldsymbol{\Sigma}_i^{\beta}\}$ and $\{\boldsymbol{\mu}_{j\to i}, \boldsymbol{\Sigma}_{j\to i}\}$, respectively. Then Eq. (S10.17) is reduced to

$$\boldsymbol{\mu}_{j\to i} = \boldsymbol{\Sigma}_{i|j}^{\mathrm{Bay}}[\boldsymbol{A}^{\mathbb{T}}\Sigma_w^{-1}\boldsymbol{\mu}_j^{\beta} + (\boldsymbol{\Sigma}_i^{\mathrm{p}})^{-1}\boldsymbol{\mu}_i^{\mathrm{p}}], \tag{S10.27}$$

$$\boldsymbol{\Sigma}_{j\to i} = \left[(\boldsymbol{\Sigma}_j^{\beta})^{-1} + \boldsymbol{\Sigma}_{i|j}^{\mathrm{Bay}}\boldsymbol{A}^{\mathbb{T}}\boldsymbol{\Sigma}_{\boldsymbol{w}}^{-1}(\boldsymbol{\Sigma}_{i|j}^{\mathrm{Bay}})^{-1}(\boldsymbol{\Sigma}_{i|j}^{\mathrm{Bay}}\boldsymbol{A}^{\mathbb{T}}\boldsymbol{\Sigma}_{\boldsymbol{w}}^{-\mathbf{1}})^{\mathbb{T}}\right]^{-1}. \tag{S10.28}$$

Let $\{\boldsymbol{\mu}_i^{\mathrm{syn}}, \boldsymbol{\Sigma}_i^{\mathrm{syn}}\}$ be the parameter of $p(x_i|\mathrm{down}(i))$, then Eq. (S10.18) becomes

$$\boldsymbol{\mu}_i^{\mathrm{syn}} = \boldsymbol{\Sigma}_i^{\mathrm{syn}}[(\boldsymbol{\Sigma}_{j\to i})^{-1}\boldsymbol{\mu}_{j\to i} + (\boldsymbol{\Sigma}_{j'\to i})^{-1}\boldsymbol{\mu}_{j'\to i} - (\boldsymbol{\Sigma}_i^{\mathrm{p}})^{-1}\boldsymbol{\mu}_i^{\mathrm{p}}], \tag{S10.29}$$

$$\boldsymbol{\Sigma}_i^{\mathrm{syn}} = [(\boldsymbol{\Sigma}_{j\to i})^{-1} + (\boldsymbol{\Sigma}_{j'\to i})^{-1} - (\boldsymbol{\Sigma}_i^{\mathrm{p}})^{-1}]^{-1}. \tag{S10.30}$$

For the cells at the end of lineages, $\boldsymbol{\mu}_i^{\mathrm{syn}} = \boldsymbol{\mu}_i^{\mathrm{p}}$ and $\boldsymbol{\Sigma}_i^{\mathrm{syn}} = \boldsymbol{\Sigma}_i^{\mathrm{p}}$. Finally, from Eq. (S10.19), we obtain

$$\boldsymbol{\mu}_i^{\beta} = \boldsymbol{\mu}_i^{\mathrm{syn}} + \boldsymbol{K}(\tau_i - \boldsymbol{C}\boldsymbol{\mu}_i^{\mathrm{syn}}), \qquad \boldsymbol{\Sigma}_i^{\beta} = \boldsymbol{\Sigma}_i^{\mathrm{syn}} - \boldsymbol{K}\boldsymbol{C}\boldsymbol{\Sigma}_i^{\mathrm{syn}}, \tag{S10.31}$$

where $\boldsymbol{K}$ is a Kalman gain defined by

$$\boldsymbol{K} = \boldsymbol{\Sigma}_i^{\mathrm{syn}}\boldsymbol{C}^{\mathbb{T}}(\boldsymbol{\Sigma}_{\boldsymbol{v}} + \boldsymbol{C}\boldsymbol{\Sigma}_i^{\mathrm{syn}}\boldsymbol{C}^{\mathbb{T}})^{-1}. \tag{S10.32}$$

This completes the recursive computation of $\bar{\beta}_i$.

Let $m$ be the label of the parent of the cell $i$. Let the parameters of $\gamma_i(\boldsymbol{x}_i)$ be $\{\boldsymbol{\mu}_i^{\mathrm{s}}, \boldsymbol{\Sigma}_i^{\mathrm{s}}\}$, then Eq. (S10.23) together with Eq. (S10.22) is represented as

$$\boldsymbol{\mu}_i^{\mathrm{s}} = \boldsymbol{\mu}_i^{\beta} + \boldsymbol{J_i}(\boldsymbol{\mu}_m^{\mathrm{s}} - \boldsymbol{A}^{\mathbb{T}}\boldsymbol{\Sigma}_{\boldsymbol{w}}^{-1}\boldsymbol{\mu}_i^{\beta}), \qquad \boldsymbol{\Sigma}_i^{\mathrm{s}} = \boldsymbol{\Sigma}_i^{\beta} + \boldsymbol{J_i}(\boldsymbol{\Sigma}_m^{\mathrm{s}} - \boldsymbol{\Sigma}_i^{\mathrm{syn}})\boldsymbol{J_i}^{\mathbb{T}}, \tag{S10.33}$$

where $\boldsymbol{J_i} = \boldsymbol{\Sigma}_i^{\mathrm{s}}\boldsymbol{A}^{\mathbb{T}}\boldsymbol{\Sigma}_{\boldsymbol{w}}^{-1}(\boldsymbol{\Sigma}_i^{\mathrm{syn}})^{-1}$. Let $\mathrm{cov}[\boldsymbol{x}_i, \boldsymbol{x}_j]$ be the covariance matrix of the state $\boldsymbol{x}_i$ and $\boldsymbol{x}_j$ under the distribution $\xi_{i,j}(\boldsymbol{x}_i, \boldsymbol{x}_j)$, that is

$$\mathrm{cov}[\boldsymbol{x}_i, \boldsymbol{x}_j] = \mathbb{E}_{\xi_{i,j}(\boldsymbol{x}_i, \boldsymbol{x}_j)}[\boldsymbol{x}_j\boldsymbol{x}_i^{\mathbb{T}}]. \tag{S10.34}$$

Then, Eq. (S10.24) implies that

$$\mathrm{cov}[\boldsymbol{x}_i, \boldsymbol{x}_j] = \boldsymbol{J_j} \boldsymbol{\Sigma}_i^{\mathrm{syn}}. \tag{S10.35}$$

This completes the recursive computation of $\gamma_i$ and $\xi_{i,j}$.

To evaluate the convergence of LEM and also conduct the model selection, the value of the likelihood $\mathcal{L}$ is required, where

$$\mathcal{L} := \mathbb{P}(\mathcal{D}). \tag{S10.36}$$

To obtain a recursive formula for $\mathcal{L}$, we define

$$\ell_i = \mathbb{P}(\mathrm{down}(i), \tau_i). \tag{S10.37}$$

For the root cell, $\mathcal{L} = \ell_{\mathrm{root}}$ holds. $\ell_i$ is calculated as

$$\ell_i = \mathbb{P}(\mathrm{down}(i), \tau_i) = \mathbb{P}(\tau_i|\mathrm{down}(i))\mathbb{P}(\mathrm{down}(i)). \tag{S10.38}$$

We calculate the first factor $g_i := \mathbb{P}(\tau_i|\mathrm{down}(i))$ in Eq. (S10.38) as

$$g_i = \int d\boldsymbol{x}_i \mathbb{P}(\tau_i|\boldsymbol{x}_i)\mathbb{P}(\boldsymbol{x}_i|\mathrm{down}(i)) = \int d\boldsymbol{x}_i \pi_{\mathrm{B}}(\tau_i|\boldsymbol{x}_i)\bar{\beta}_i(\boldsymbol{x}_i). \tag{S10.39}$$

We next calculate the second factor $\mathbb{P}(\mathrm{down}(i))$ in Eq. (S10.38). By integrating the first line of (S10.12) with respect to $\boldsymbol{x}_i$, we obtain

$$1 = \frac{\mathbb{P}(\mathrm{down}(j), \tau_j)\mathbb{P}(\mathrm{down}(j'), \tau_{j'})}{\mathbb{P}(\mathrm{down}(i))} \int d\boldsymbol{x}_i \, \bar{\beta}_{j\to i}(\boldsymbol{x}_i)\bar{\beta}_{j'\to i}(\boldsymbol{x}_i)\mathbb{P}(\boldsymbol{x}_i)^{-1}. \tag{S10.40}$$

Hence, by letting

$$h_i := \bar{\beta}_{j\to i}(\boldsymbol{x}_i)\bar{\beta}_{j'\to i}(\boldsymbol{x}_i)\mathbb{P}(\boldsymbol{x}_i)^{-1}, \tag{S10.41}$$

we obtain

$$\mathbb{P}(\mathrm{down}(i)) = \ell_j \ell_{j'} h_i. \tag{S10.42}$$

In summary, the likelihood is recursively computed as

$$\ell_i = \ell_j \ell_{j'} g_i h_i, \tag{S10.43}$$

where $g_i$ and $h_i$ are given by Eqs. (S10.39) and (S10.41), respectively. At the end of the lineage in the tree, $\ell_i$ is initialized as $\ell_i = \int d\boldsymbol{x}_i \pi_{\mathrm{B}}(\tau_i|\boldsymbol{x}_i)\mathbb{P}(\boldsymbol{x}_i)$. At the root cell, we obtain $\mathcal{L} = \ell_{\mathrm{root}}$.

By a direct calculation, the calculation of the factors $g_i$ and $h_i$ are reduced to

$$g_i = \mathrm{Normal}(\tau_i|\boldsymbol{C}\boldsymbol{\mu}_i^{\mathrm{syn}}, \boldsymbol{C}\boldsymbol{\Sigma}_i^{\mathrm{syn}}\boldsymbol{C}^{\mathbb{T}} + \Sigma_v), \tag{S10.44}$$

where $\mathrm{Normal}(\boldsymbol{x}|\boldsymbol{\mu}, \boldsymbol{\Sigma})$ is the value of the probability density function of normal distribution with mean $\boldsymbol{\mu}$ and covariance matrix $\boldsymbol{\sigma}$ the at $\boldsymbol{x}$, and

$$h_i = \sqrt{\frac{|\boldsymbol{\Sigma}_i^{\mathrm{p}}||\boldsymbol{\Sigma}_i^{\mathrm{syn}}|}{|\boldsymbol{\Sigma}_{j\to i}||\boldsymbol{\Sigma}_{j'\to i}|}} \exp\left[\frac{1}{2}\Big(((\boldsymbol{\mu}_i^{\mathrm{syn}})^{\mathbb{T}}(\boldsymbol{\Sigma}_i^{\mathrm{syn}})^{-1}\boldsymbol{\mu}_i^{\mathrm{syn}} + (\boldsymbol{\mu}_i^{\mathrm{p}})^{\mathbb{T}}(\boldsymbol{\Sigma}_i^{\mathrm{p}})^{-1}\boldsymbol{\mu}_i^{\mathrm{p}} \right.$$
$$\left. -(\boldsymbol{\mu}_{j\to i})^{\mathbb{T}}(\boldsymbol{\Sigma}_{j\to i})^{-1}\boldsymbol{\mu}_{j\to i} - (\boldsymbol{\mu}_{j'\to i})^{\mathbb{T}}(\boldsymbol{\Sigma}_{j'\to i})^{-1}\boldsymbol{\mu}_{j'\to i}\Big)\right]. \tag{S10.45}$$

For the cells at the end of lineage in the tree, $l_i$s are initialized as

$$l_i = \mathrm{Normal}(\tau_i|\boldsymbol{C}\boldsymbol{\mu}_i^{\mathrm{p}}, \boldsymbol{C}\boldsymbol{\Sigma}_i^{\mathrm{p}}\boldsymbol{C}^{\mathbb{T}} + \Sigma_v). \tag{S10.46}$$

### S11 Details on numerical experiments

We implemented all algorithms in C++14. In the E-step of LEM, we used a scaling technique to avoid an underflow problem. In the recursive computation of $\alpha_i(\boldsymbol{x})$ and $\beta_i(\boldsymbol{x})$, they are normalized to 1 at each step. This normalization does not affect the computed $\gamma_i(\boldsymbol{x}_i)$ and $\xi_{ij}(\boldsymbol{x}_i, \boldsymbol{x}_j)$ owing to the fact that the recursive formula is in the bilinear form of $\alpha_i(\boldsymbol{x})$ and $\beta_i(\boldsymbol{x})$. The probability density function of the gamma distribution was computed by a numerical differentiation of an incomplete gamma function. This enables us to compute the probatiblity density function of the gamma distribution with a large shape parameter. Incomplete gamma functions were computed by Boost C++ library 1.64.0. In the M-step, the modified moment-maching equation of the gamma distribution was solved numerically by the Newton method [23]. The parameters for a continuous state model were updated by using formulae in Ref. [21]. We used Eigen 3.3.3 for matrix calculations. The division times of the leaf cells are unknown due to the truncation of the experiment. In the discrete-state model, we treated such division times as unobserved variables in the the BW algorithm. In the E-step of the BW algorithm, we computed the posterior distributions of such division times conditioned on the event that such division times are longer than the observed ones. Then, in the M-step, we updated the parameters using such posterior distributions. In a continuous state model, we just ignored such cells for simplicity. The figures were ploted either by Python 3.6.1 or by Mathematica.

We executed our LEM in the following environment. The operating system is Windows 10 Education version 1803. The CPU is Intel Core i7-6820HQ 2.70 GHz. The random access memory is 16GB. We used no parallel computation and GPUs. The source code was complied by Visual Studio Community 2017 version 15.5.2 in the Release mode.

### S12 Details on the estimation of the synthetic discrete model

We here describe the detail of the synthetic lineage tree generated from the discrete-state model. The states of the model are denoted as $\Omega = \{f, s\}$. The symbol $f$ and $s$ represent the fast and slow growing states, respectively. The distribution of division times follows the gamma distribution with the shape parameter $a$ and the scaling parameter $b$. The parameters of the fast growing state are $a_f = 20$ and $b_f = 1$, and those of the slow growing state are $a_s = 30$ and $b_s = 1$. The stochastic transition matrix of the state switching is set to be $\mathbb{T}_{\mathrm{F}}(f|f) = \mathbb{T}_{\mathrm{F}}(s|s) = 0.7$ and $\mathbb{T}_{\mathrm{F}}(f|s) = \mathbb{T}_{\mathrm{F}}(s|f) = 0.3$.

We generated the synthetic lineage tree based on this model as follows. The tree was rooted by a cell at time 0 with age 0, and a population of the cells is derived based on the model until $t = 220$. The state of the root cell was sampled from the stationary distribution of a population following the model. To obtain a sample from the stationary distribution and the growth rate $\lambda$, a synthetic lineage was generated beforehand starting from a root cell with state $f$ up to $t = 500$. This procedure is similar to the preincubation of a population before starting observation of a lineage tree in an experiment.

Upon division of a cell, the division times and the states of its daughters were sampled on the basis of the above gamma distribution and the stochastic transition matrix. We recursively conducted this procedure to obtain the lineage tree. The cells that were born after $t = 220$ were ignored. The sampling was computed by C++ standard library, and a generator of pseudorandom numbers mt19937_64. The seed of the generator was random_device.

In the procedure of the inference, we chose the initial value of the parameters as follows. We first compute the maximum likelihood estimator for the division times under the assumption that there is no state. The initial values of the parameters for the division times are set to this maximum likelihood estimator. The initial value of the estimated transition matrix was set to be the uniform distribution, $\mathbb{T}_{\mathrm{F}}(\boldsymbol{x}|\boldsymbol{x}') = 1/|\Omega|$.

In the inference, $\pi_{\mathrm{B}}(\tau|\boldsymbol{x})$ rather than $\pi_{\mathrm{F}}(\tau|\boldsymbol{x})$ is estimated. If $\pi_{\mathrm{F}}(\tau|\boldsymbol{x})$ is a gamma distribution as we assumed, $\pi_{\mathrm{B}}(\tau|\boldsymbol{x})$ becomes also a gamma distribution, because of the relation between the forward and retrospective processes, $\pi_{\mathrm{B}}(\tau|\boldsymbol{x}) \propto e^{-\lambda\tau}\pi_{\mathrm{F}}(\tau|\boldsymbol{x})$ (See Eq. [3] in the main text). Thereby, we can use the gamma distribution even in the procedure of the inference. The parameters $a_{\mathrm{B}}$ and $b_{\mathrm{B}}$ of $\pi_{\mathrm{B}}(\tau|\boldsymbol{x})$ for a fixed $\boldsymbol{x}$ can be related with those of $\pi_{\mathrm{F}}(\tau|\boldsymbol{x})$, $a_{\mathrm{F}}$ and $b_{\mathrm{F}}$, as $a_{\mathrm{B}} = a_{\mathrm{F}}$ and $b_{\mathrm{B}} = b_{\mathrm{F}} + \lambda$. By using these formulae, we can obtain the parameters of $\pi_{\mathrm{F}}(\tau|\boldsymbol{x})$.

To estimate the statistical variations of the estimated parameters, we conducted independent estimations 1000 times with the same true parameters.

Table S12.1: The summary statistics of the estimated parameters of the first model obtained by 1000 times executions of our algorithm. The means of the estimated values are close to the true values and the standard deviations of them are small. After the correction of the survivor bias, the estimated value of $b$ is improved.

| parameter | true value | mean | standard deviation | corrected mean | corrected standard deviation |
|---|---|---|---|---|---|
| $\mathbb{T}_{\mathrm{F}}(f\|f)$ | 0.7 | 0.698 | 0.0259 | — | — |
| $\mathbb{T}_{\mathrm{F}}(f\|s)$ | 0.3 | 0.303 | 0.0351 | — | — |
| $a_f$ | 20 | 20.8 | 2.42 | 20.8 | 2.42 |
| $b_f$ | 1 | 1.04 | 0.129 | 1.01 | 0.129 |
| $a_s$ | 30 | 32.4 | 5.90 | 32.4 | 5.90 |
| $b_s$ | 1 | 1.08 | 0.190 | 1.06 | 0.190 |

Table S12.2: The summary statistics of the estimated parameters of the second model obtained by 100 times executions of our algorithm. We excluded the five times of execution in which the estimated parameters diverge or are far from the true ones. Since the survivor bias does not distort the estimation of $\mathbb{T}_{\mathrm{F}}$, the corrected mean and the corrected standard deviation of $\mathbb{T}_{\mathrm{F}}$ is left blank. After the correction of the survivor bias, the estimated value of $b$ is correctly recovered.

| parameter | true value | mean | standard deviation | corrected mean | corrected standard deviation |
|---|---|---|---|---|---|
| $\mathbb{T}_{\mathrm{F}}(f\|f)$ | 0.8 | 0.732 | 0.0172 | — | — |
| $\mathbb{T}_{\mathrm{F}}(f\|s)$ | 0.2 | 0.180 | 0.0153 | — | — |
| $a_f$ | 1.5 | 1.48 | 0.640 | 1.48 | 0.640 |
| $b_f$ | 1.0 | 1.531 | 0.0884 | 0.977 | 0.0884 |
| $a_s$ | 450 | 421 | 42.4 | 421 | 42.4 |
| $b_s$ | 300 | 282 | 28.4 | 281 | 28.4 |

The summary statistics of the estimated parameters are shown in Table S12.1. Since this transformation of the parameters from $\pi_{\mathrm{B}}$ to $\pi_{\mathrm{F}}$ is linear, only the means are transformed.

Since the survivor bias in the model was small, we conducted estimation on another model with different parameter values. The parameters of the fast growing state are $a_f = 1.5$ and $b_f = 1$, and those of the slow growing state are $a_s = 450$ and $b_s = 300$. The stochastic transition matrix of the state switching is set to be $\mathbb{T}_{\mathrm{F}}(f|f) = \mathbb{T}_{\mathrm{F}}(s|s) = 0.8$ and $\mathbb{T}_{\mathrm{F}}(f|s) = \mathbb{T}_{\mathrm{F}}(s|f) = 0.2$. The tree was rooted by a cell at time 0 with age 0, and a population of the cells is derived on the basis of the model until $t = 15$. We conducted independent estimations 100 times with the same true parameters. The estimated parameter diverges in two of the 100 trials and the estimated parameters were far from the true ones in three of the 100 trials. The summary statistics of the estimated parameters are shown in Table S12.2.

In Figure 5 (g) and (h) in the main text, we plotted the estimated $\pi_{\mathrm{B}}(\tau|f)$ and $\pi_{\mathrm{B}}(\tau|s)$ weighted by the stationary distributions of the states. The histogram of the division times was calculated from all the cells but at the leaves in the tree.

### S13 Details on the estimation of the continuous model

We randomly chose the initial values of the parameters used in the algorithm as follows. Because the covariance matrices $\boldsymbol{\Sigma_w}$ and $\Sigma_v$ are symmetric and positive definite, we need the following special technique: to construct a $k \times k$ symmetric positive definite matrix, we first generated random vectors $v_i$ $(i = 1, 2, \ldots, k)$, each element of which follows the uniform distribution on $[-5, 5]$. Then, we obtained $\Sigma$ as $\Sigma = \sum_{i=1}^{k} v_i {v_i}^{\mathbb{T}}$. For the model considered here, we chose $k = 2$ for $\boldsymbol{\Sigma_w}$ and $k = 1$ for $\Sigma_v$. For the other parameters, $\boldsymbol{A}$ and $\boldsymbol{C}$, each element was uniformly sampled from $[-5, 5]$. Finally, we ran the estimation algorithm for 1000 different initial values and adopted the parameters with the largest likelihood.

To compare the estimated parameters with the true parameters correctly, we need to transform the coordinate of the estimated parameters into that of the true parameters, because a different set of parameters might represent the same model. Specificially, by transforming the coordinate of the state $\boldsymbol{x}$ into $\boldsymbol{Lx}$ by any $k \times k$ nonsingular matrix $\boldsymbol{L}$, we can obtain a different set of parameters

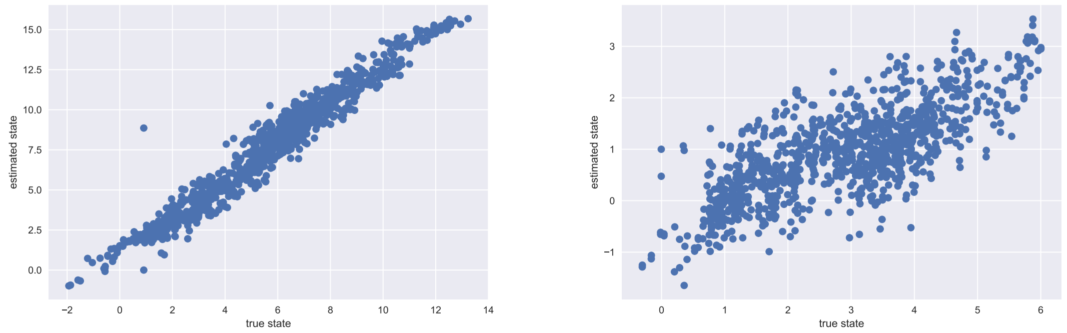


Figure S14.1: Comparison between the true state and the expectation of the estimated state by LEM. The true state and the expectation of the estimated state is plotted for each cells in the lineage tree. The left figure shows $x_1$ and the right figure shows $x_2$. The horizontal axis is the true state and the vertical axis is the expectation of the estimated state.

that represent the same model as

$$\begin{aligned} \boldsymbol{A}' &= \boldsymbol{L}\boldsymbol{A}\boldsymbol{L}^{-1}, \quad \boldsymbol{C}' = \boldsymbol{C}\boldsymbol{L}^{-1}, \\ \boldsymbol{\Sigma}'_{\boldsymbol{w}} &= \boldsymbol{L}\boldsymbol{\Sigma}_{\boldsymbol{w}}\boldsymbol{L}^{\mathbb{T}}, \quad \Sigma'_v = \Sigma_v. \end{aligned} \tag{S13.1}$$

To remove this ambituity in the coordinate, we transformed the estimated parameters as follows. First, we diagonalized the estimated $\boldsymbol{\Sigma}_{\boldsymbol{w}}$ as $\boldsymbol{\Sigma}'_{\boldsymbol{w}} = \boldsymbol{L}\boldsymbol{\Sigma}_{\boldsymbol{w}}(\boldsymbol{L})^{\mathbb{T}}$. Here, the transformation matrix $\boldsymbol{L}$ was determined up to constant multiplications to each eigenvector in $\boldsymbol{L}$ and the order of the eigenvectors. We next chose these constant multiplications so that $\boldsymbol{C}' = \boldsymbol{C}\boldsymbol{L}^{-1} = (1, 1)$. We finally chose the order of the eigenvalues so that the coordinate of the transformed parameters by Eq. (S13.1), i.e. $\boldsymbol{A}', \boldsymbol{C}', \boldsymbol{\Sigma}'_{\boldsymbol{w}}$, and $\Sigma'_v$, became the closest to that of the true parameters. Specifically, we multiplied the transformation matrix $\boldsymbol{L}$ by

$$\begin{pmatrix} 0 & 1 \\ 1 & 0 \end{pmatrix} \tag{S13.2}$$

from the left if the $(0, 1)$-element of $\boldsymbol{A}'$ is smaller than the $(1, 0)$-element of $\boldsymbol{A}'$. In summary, by the above transformations, we obtained parameters $\boldsymbol{A}', \boldsymbol{C}', \boldsymbol{\Sigma}'_{\boldsymbol{w}}$, and $\Sigma'_v$, the coordinate of which may coincides with that of the true parameters.

### S14 Comparison between the true latent state and the mean of the estimated state in the contionous model

We validate that our LEM reconstructs the state of the cells in the continuous model. To this end, we compared the true state of a synthetic lineage tree and the estimated states. We generated synthetic lineage trees by using almost the same model as Eq. [14] in the main text. Here, we used larger noise $\Sigma_{\boldsymbol{w}}$ than [14] in the main text, defined by

$$\Sigma_{\boldsymbol{w}} = \begin{pmatrix} 0.1 & 0 \\ 0 & 0.1 \end{pmatrix}, \tag{S14.1}$$

since our interest is whether the estimated states can follow the true state. The latent states moves more broadly in this model and we can check whether the estimated state by LEM follows the true latent state. We used the coordinate transformation Eq. (S13.1) before the M-step of the LEM to compare the latent states correctly. The comparison between the true state and the expectation of the estimated state is shown in Figure S14.1. We can see that the estimated states follow the true state. The correlation coefficients between the true state and the expectation of the estimated state are 0.979 for the first component $x_1$ and 0.750 for the second component $x_2$. This result quantitatively verifies that our LEM can reconstructs the state of the cells.

#### S15 An application of *E. coli* cells lineage trees

We next inferred the latent states of the *E. coli* cells in the lineage trees observed by using the dynamics cytometer in Hashimoto *et al.* [24] (Fig. 2 (d) in the main text). The population of *E. coli* (F3 rpsL-gfp strain) was observed every one minute in the M9 minimum medium supplemented with 0.2% glucose at 37°C. We first applied LEM for the discrete model and determined the number of the latent states by Akaike Information Criteria (AIC) [25]. The best number of the discrete states was estimated to be 1, which means that the discrete model with no latent state fits the data the best (data not shown). However, this result cannot explain the non-zero correlation ($r = 0.2082$) between the division times of the mother-daughter pair observed in our data set. Here, we ensured that the correlation is significant by employing the standard deviation of the correlation coefficient estimated by 1000 times bootstrapping. The estimated standard deviation was 0.0333, which is smaller than $r = 0.2082$. A potential reason why the discrete model could not capture this correlation and the associated latent states may be that the latent states are not distinct enough to be detected by the discrete-state model. This hypothesis suggests that the latent state is better represented by the continuous rather than the discrete model. To validate that, we applied LEM of the continuous-state model, in which the dimension $k$ of the state space $\Omega = \mathbb{R}^k$ was again determined by AIC. Then, we found $k = 3$ to be the dimension of the best continuous model. This continuous model is better than any disrete model in terms of AIC. The AIC of the best continous model ($k = 3$) is -307.415, while that of the best discrete model ($k = 1$) is -671.030. To compare these two AIC values , we need to be careful about the coordinate-dependency of the AIC. Since the log-likelihood and the AIC depend on the coordinate, we must calculate the AIC of the both models in the same coordinate. However, while the discrete model used the scale $\tau$ without any transformation, the continous model used the logarithmic scale $\log \tau$. Therefore, we calculated the log-likelihood and the AIC of the discrete model in the logarithmic scale.

Of the three components of the inferred latent state of the continous model, the first one has the fastest time-scale of approximately one generation, whereas the third one changes slowly over generations (Fig. 8 in the main text). We also obtained the inferred dynamics of the latent state $\boldsymbol{x}$ over the lineage tree as in Fig. 8 in the main text and its parameter values as follows:

$$\begin{aligned} \boldsymbol{A} &= \begin{pmatrix} -0.731 & 0.438 & 0.032 \\ -2.51 & 1.124 & 0.062 \\ -0.262 & 0.0068 & 1.007 \end{pmatrix}, \quad & \boldsymbol{C} &= \begin{pmatrix} 1 & 1 & 1 \end{pmatrix}, \\ \boldsymbol{\Sigma}_w &= \begin{pmatrix} 0.055 & 0 & 0 \\ 0 & 0.038 & 0 \\ 0 & 0 & 0.016 \end{pmatrix}, \quad & \Sigma_v &= 0.04. \end{aligned} \tag{S15.1}$$

In the data of Hashimoto *et al.* [24], the cells exceeding the capacity of the chamber were flown out, and therefore we can no longer observe the flown cells. In particular, we cannot know the division time of them. In our estimation, the flown cells were excluded from the data, which corresponding to pruning the leaves in the lineage tree. For the initial values of the parameters in LEM, we chose them in the same way as the previous section. We ran the estimation algorithm for 1000 different initial values and adopted the parameters with the largest likelihood, then we obtained the estimated parameters in Eq. (S15.1).

As shown in Fig. S15.1 (a), the likelihood increases monotonically in terms of $k$, and $k = 3$ is the dimension above which the likelihood starts saturating, indicating that LEM convergences and the inference is achieved appropriately. In order to validate the significance of $k = 3$, we firstly simulated the continuous-state model without latent state ($k = 0$) for two parameter sets to obtain synthetically lineage tree data, and then applied LEM to infer the dimensionality from the synthetic data. For all 100 independent simulations and the subsequent inferences, we have obtained $k = 0$ as the inferred dimensionality (data not shown), demonstrating that LEM rarely detect a wrong latent state if it does not exist. To check the validity further, we also conducted a bootstrap analysis in which we generated surrogate trees by randomly swapping the division times of the cells in the *E. coli* lineage tree (Fig. 2 (d) in the main text) and applied LEM to the surrogates. Because the division times of the cells in the surrogate trees can be approximated to be mutually independent due to the random swapping, the surrogate trees can effectively work as the data from the null hypothesis of no latent state. Of 100 trials, $k = 0$ was inferred in most of cases (Fig. S15.1 (b)). In the rest of the trials, $k = 1$ and $k = 4$ were obtained. All the trials with $k = 4$ inferred are accompanied by much higher likelihoods than the case of $k = 0$ ((Fig. S15.1 (a)) and

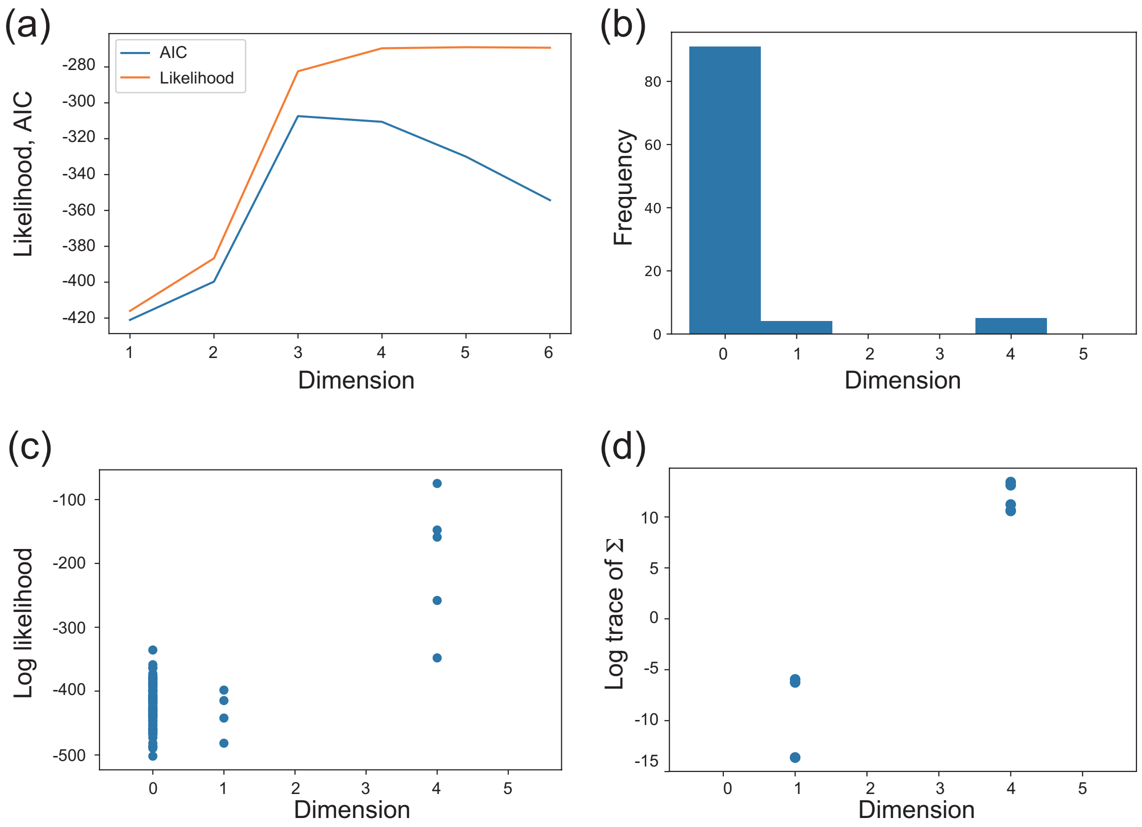


Figure S15.1: (a) The log-likelihood and AIC as functions of the dimension $k$. (b) Histogram of AICs obtained by the bootstrap analysis using 100 independently surrogated trees. (c, d) A scatter plots of the likelihoods (c) and the variances of the latent state $\boldsymbol{\Sigma}_w$ (d) from the surrogated trees. Each point correspond to the value obtained from each surrogated tree.

irregularly large variances for the latent state $\boldsymbol{w}$ (Fig. S15.1 (d)). Such large variances effectively allow the latent state to arbitrarily fit to the observations. Therefore, $k = 4$ is probably due to an inappropriate convergence of the EM algorithm, which has also been reported to occur when it is applied to the MLE of models with latent states [21]. In contrast, the results of $k = 1$ show the likelihood and the variance, comparable to those of $k = 0$. This suggests that the model with $k = 0$ sometime generates samples being similar to those from $k = 1$. However, the lack of $k = 2$ indicates that the probability to obtain $k > 1$ from the model of $k = 0$ by chance is much less than 1/100. Lastly, we also applied LEM to another *E. coli* tree and obtained $k = 3$ for this data set (data not shown). Therefore, $k = 3$ inferred from Fig. 2 (d) in the main text should have a sufficient statistical significance.

Next, we investigated how the latent state represents the stochastic behavior of the division times. From the assumption of the continuous model, the posterior average of the division time of the cell $i$ in the tree is obtained as

$$\langle \log \tau_i \rangle = \boldsymbol{C}\boldsymbol{x}_i = x_i^1 + x_i^2 + x_i^3. \tag{S15.2}$$

The comparison of $\langle \log \tau_i \rangle$ with the actual observation of the division time $\log \tau_i$ shows that the intercellular variation of the division times is mainly accounted by the fluctuation of the latent state (Fig. S15.2 (a)), which is also reflected in the small value of the state-independent fluctuation $\Sigma_v$ (Eq. (S15.1)). The dissection of $\langle \log \tau_i \rangle$ into each component of the latent state also indicates that $x^1$ and $x^2$ mainly represent the fluctuation of the division time whereas $x^3$ encodes its average value (Fig. S15.2 (a)).

Then, we also analyzed how the latent state conveys the information on the division statistics over generations. By using $\boldsymbol{A}$ and $\boldsymbol{x}$ inferred, we can predict the division time of the daughter cells from the latent states of their mothers as

$$\langle \log \tau_{i+1} \rangle = \boldsymbol{C}\boldsymbol{A}\boldsymbol{x}_i. \tag{S15.3}$$

where we abuse the notation $i+1$ to mean the label of a daughter cell of the cell $i$. Similarly, we can predict the division times of the grand daughter cells. As shown in Fig. S15.2 (c), the latent state effectively captures the relationship of the division times over generation, and thereby, the posterior averages of the division times, $\langle \log \tau_i \rangle$ and $\langle \log \tau_{i+1} \rangle$, also reproduce the correlation between the mother-daughter pairs as $r = 0.2034$ (Fig. S15.2 (b)).

Finally, we clarify how the inter-generation information is encoded in the latent state and its dynamics by plotting the phase space dynamics of the latent state (Fig. S15.2 (d)). The latent dynamics had fast and slow components: the fast one is basically the projective dynamics to an one-dimensional sub-manifold in the $x^1$-$x^2$ plane(Fig. S15.2 (e)), whereas the slow one is a dynamics formed in the the sub-manifold and $x^3$(Fig. S15.2 (d)). This result demonstrates that $x^3$ is not only encoding the average value of the division time, but also the information of the division times of its descendants. In conclusion, the slow dynamics suggests an existence of a slow regulatory factor underlying the noisy behavior of the division times and being inherited over generations.

In conclusion, we have identified the latent low-dimensional states of the cells, which are inherited over a couple of generations at least. The inferred states successfully capture the underlying effective inheritance dynamics of the division times over generations even though the correlation of the observed division times between the mother-daughter pairs is subtle presumably because of the stochastic nature of the cellular replication.

### References


[1] Jelinek F (1997) *Statistical methods for speech recognition.* (MIT press).

[2] Rabiner LR, Juang BH (1993) *Fundamentals of speech recognition.* (PTR Prentice Hall Englewood Cliffs) Vol. 14.

[3] Manning CD, Schütze H (1999) *Foundations of statistical natural language processing.* (MIT press).

[4] Wang P, et al. (2010) Robust growth of escherichia coli. *Current Biology* 20(12):1099 – 1103.

[5] Hormoz S, Desprat N, Shraiman BI (2015) Inferring epigenetic dynamics from kin correlations. *Proceedings of the National Academy of Sciences* 112(18):E2281–E2289.

[6] Hormoz S, et al. (2016) Inferring cell-state transition dynamics from lineage trees and endpoint single-cell measurements. *Cell Systems* 3(5):419 – 433.e8.

[7] Hicks DG, Speed TP, Yassin M, Russell SM (2018) Statistical inference in cell lineage trees. *bioRxiv.*

[8] Hicks DG, Speed TP, Yassin M, Russell SM (2018) Maps of variability in cell lineage trees. *bioRxiv.*

[9] Phillips NE, Mandic A, Omidi S, Naef F, Suter DM (2019) Memory and relatedness of transcriptional activity in mammalian cell lineages. *Nature Communications* 10(1):1208.

[10] Olariu V, et al. (2009) Modified variational bayes em estimation of hidden markov tree model of cell lineages. *Bioinformatics* 25(21):2824–2830.

[11] Failmezger H, et al. (2018) Clustering of samples with a tree-shaped dependence structure, with an application to microscopic time lapse imaging. *Bioinformatics* p. bty939.

[12] Kuzmanovska I, Milias-Argeitis A, Mikelson J, Zechner C, Khammash M (2017) Parameter inference for stochastic single-cell dynamics from lineage tree data. *BMC Systems Biology* 11(1):52.

[13] Kuchen EE, Becker N, Claudino N, Hofer T (2018) Long-range memory of growth and cycle progression correlates cell cycles in lineage trees. *bioRxiv.*

[14] Sughiyama Y, Nakashima S, Kobayashi TJ (2018) Fitness response relation of a multi-type age-structured population dynamics. *ArXiv e-prints.*

[15] Marc H, Adélaïde O (2016) Nonparametric estimation of the division rate of an age dependent branching process. *Stochastic Processes and their Applications* 126(5):1433 – 1471.

[16] Marguet A (2016) Uniform sampling in a structured branching population. *ArXiv e-prints.*

[17] Wang H, Qian H (2007) On detailed balance and reversibility of semi-markov processes and single-molecule enzyme kinetics. *Journal of Mathematical Physics* 48(1):013303.

[18] Maes C, Netočný K, Wynants B (2009) Dynamical fluctuations for semi-markov processes. *Journal of Physics A: Mathematical and Theoretical* 42(36):365002.

[19] Fort G, Roberts GO (2005) Subgeometric ergodicity of strong markov processes. *Ann. Appl. Probab.* 15(2):1565–1589.

[20] Lancaster P (1964) On eigenvalues of matrices dependent on a parameter. *Numer. Math.* 6(1):377–387.

[21] Christopher B (2006) *Pattern Recognition and Machine Learning.* (Springer-Verlag New York).

[22] Laferte JM, Perez P, Heitz F (2000) Discrete markov image modeling and inference on the quadtree. *IEEE Transactions on Image Processing* 9(3):390–404.

[23] Kanno Y, Tsuchiya T (2014) *Optimization and Variational Methods (in Japanese).* (Maruzen Publishing Tokyo).

[24] Hashimoto M, et al. (2016) Noise-driven growth rate gain in clonal cellular populations. *Proceedings of the National Academy of Sciences* 113(12):3251–3256.

[25] Akaike H (1998) *Information Theory and an Extension of the Maximum Likelihood Principle*, eds. Parzen E, Tanabe K, Kitagawa G. (Springer New York, New York, NY), pp. 199–213.

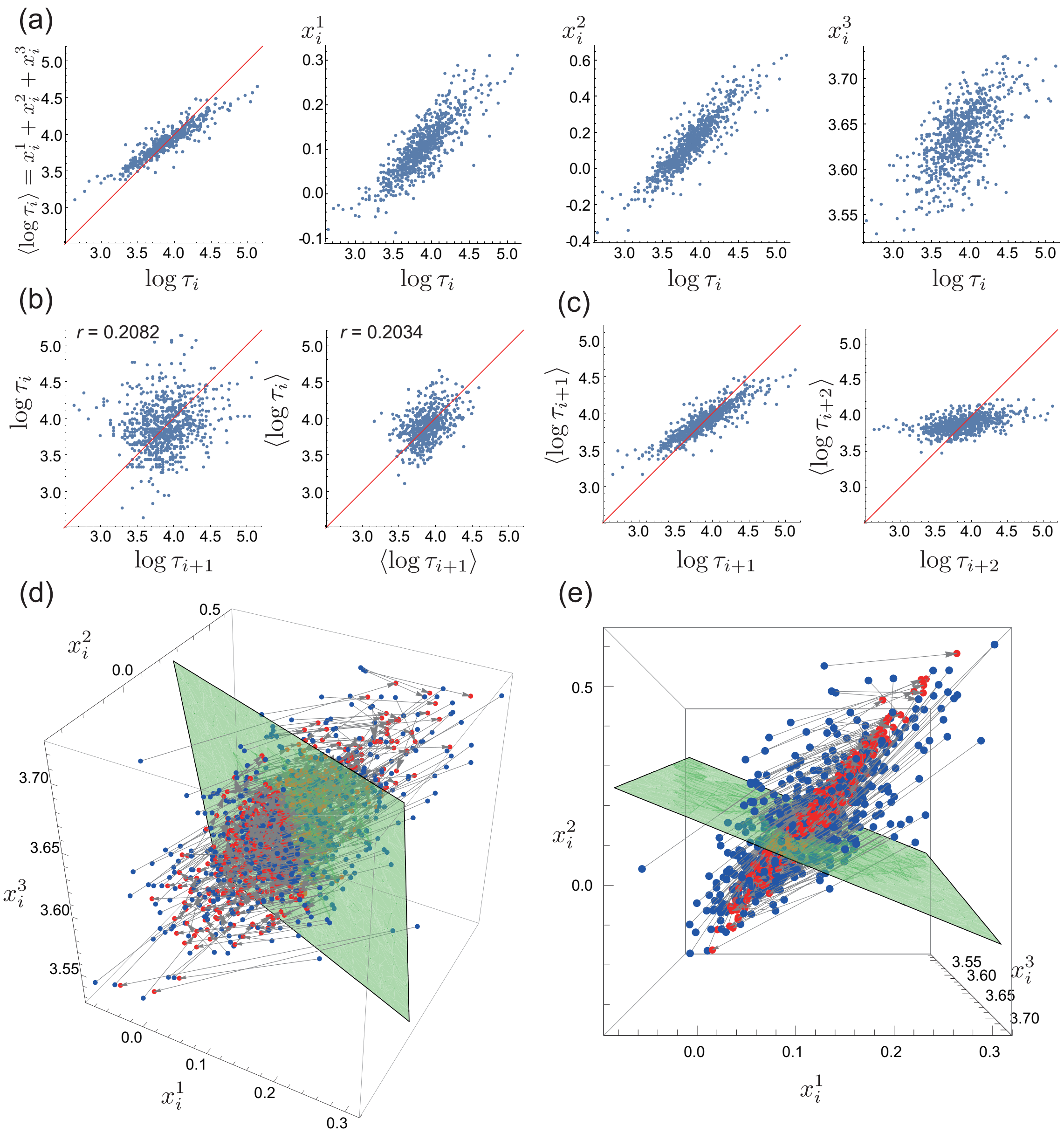


Figure S15.2: (a) Comparisons of the actual division times, $\log \tau_i$, with the predicted division times, $\langle \log \tau_i \rangle$, and with the three components of $\boldsymbol{x}_i$. Each point corresponds to each cell in the tree. (b) Comparisons of the mother's and its daughter's division times by using the actual observations (left) and the predicted values (right). Each point corresponds to each mother-daughter pair. (c) Comparisons of the actual division times of the daughter (left, $\log \tau_{i+1}$) and grand-daughter (right, $\log \tau_{i+2}$) cells with their predicted values $\langle \log \tau_{i+1} \rangle$ and $\langle \log \tau_{i+2} \rangle$ obtained from the latent states of the corresponding mother cells $\boldsymbol{x}_i$. (d) State-space representation of the dynamics of the latent state. Each blue point represents the latent state of each cell $\boldsymbol{x}_i$, and the corresponding red point connected by the gray arrow is its mapped state $A\boldsymbol{x}_i$. The green plane is an instance of the surface satisfying $x^1 + x^2 + x^3 = \text{const.}$. A subset of cells that are on the same $x^1 + x^2 + x^3 = \text{const.}$ plane generates the same predicted value of the division time. The green plane in the plot is obtained for $\text{const.} = \tau_{\text{av}}$ where $\tau_{\text{av}}$ is the sample average of the division times. (e) The same 3D plot as (d) but rendered from the top-view.
